# PR+ progenitors contribute to all mammary epithelial lineages

**DOI:** 10.1101/2025.04.16.648786

**Authors:** Pirashaanthy Tharmapalan, Matthew Waas, Curtis W McCloskey, Virginie Defamie, Kazeera Aliar, Bruno M Simoes, Robert B Clarke, Sacha J Howell, Thomas Kislinger, Hal K Berman, Paul Waterhouse, Rama Khokha

## Abstract

The central role of progesterone in breast biology and cancer is undeniable. Progesterone is a potent mitogen for mammary stem cell expansion and essential for mammopoiesis, yet only a fraction of the luminal epithelium is described to express the progesterone receptor (PR). Progestins in contraceptives and HRT regimens increases breast cancer risk, whilst progesterone inhibition reduces mammary tumorigenesis. Understanding PR within mammary epithelial dynamics is imperative, especially given mammary stem/progenitors cells are the cells-of-origin in breast cancer. Here, we show PR-primed progenitors contribute to both mammary lineages and expose a novel PR+ basal population, challenging current dogma in the field. PR lineage-tracing yields an unprecedented contribution to the luminal and basal compartments. We uncover an asymmetrically dividing PR-primed subpopulation, which has unique bipotent clonogenic capacity and forms TEB-like outgrowths upon transplantation. We enumerate PR+ basal cells and dissect their proteomic landscape, establishing PR+ basal cells as a discrete basal subset, disparate from luminal PR+ cells. Finally, we identify PR+ basal cells in multiple proteomes, scRNAseq datasets and localize PR in clinical breast specimens. This forms a new foundation for comprehending hormone receptor patterning in the breast, not only based on lineage-identity but also the progenitor-progeny hierarchy. Our study shifts the current paradigm of breast biology with implications for breast cancer treatment given the growing interest in anti-progestin based primary prevention strategies.

## INTRODUCTION

As a key hormone essential for normal mammopoiesis, progesterone plays a demonstrated role in tertiary branching and lobuloalveologenesis during pubertal development and pregnancy^1–5^. Progesterone activates breast stem cells during each reproductive cycle in the adult female, eliciting notable cell proliferation during the progesterone-high luteal phase in contrast to the resting oestrogen-high follicular phase^6–10^. Estrogen receptor (ER) or progesterone receptor (PR) positive cells are typically found in the luminal layer of mammary epithelium whilst paracrine effectors mediate hormonal cues to the basal compartment that harbors mammary stem cells (MaSC)^11–14^. Progesterone is implicated in breast cancer^15–17^, and along with ER and HER2, the PR status of breast tumors is routinely used in clinical diagnostics. Increased exposure to progestins in contraceptives or hormone replacement therapy regimens (HRT)^18,19^ increases breast cancer incidence whereas estrogen alone HRT may even be protective^20^. Inhibition of PR signalling is well known to reduce mammary tumorigenesis^21–25^. Thus, dissecting the unique role of PR beyond ER is vital, both within the normal mammary epithelial hierarchy and in instructing mammary stem/progenitor dynamics, the postulated cells-of-origin for breast cancer.

## RESULTS

### Lineage tracing exposes unexpected PR-primed progenitors

To delineate the fate of PR+ cells in the mammary epithelial hierarchy, we generated a constitutive *PR^Cre^ Rosa26^mTmG^*lineage tracing model. In these mice, PR+ cells and their progeny are indelibly eGFP labeled following Cre-mediated excision of tdTomato in PR-expressing cells (designated as GFP+, Fig. 1A). *In situ* analysis of adult virgin murine mammary glands revealed the vast majority of epithelial cells were GFP+, in both the luminal and basal lineages (Fig. 1B). This unexpected widespread GFP labelling following PR-lineage tracing diverges from the reported ER-lineage tracing study where ER-primed cells were detected only in a luminal subset^26^. tdTomato+ PR naïve epithelial cells (designated hereafter as TD+) were sparse, scattered sporadically or seen as rare cell clusters within larger ducts.

**Figure 1.**
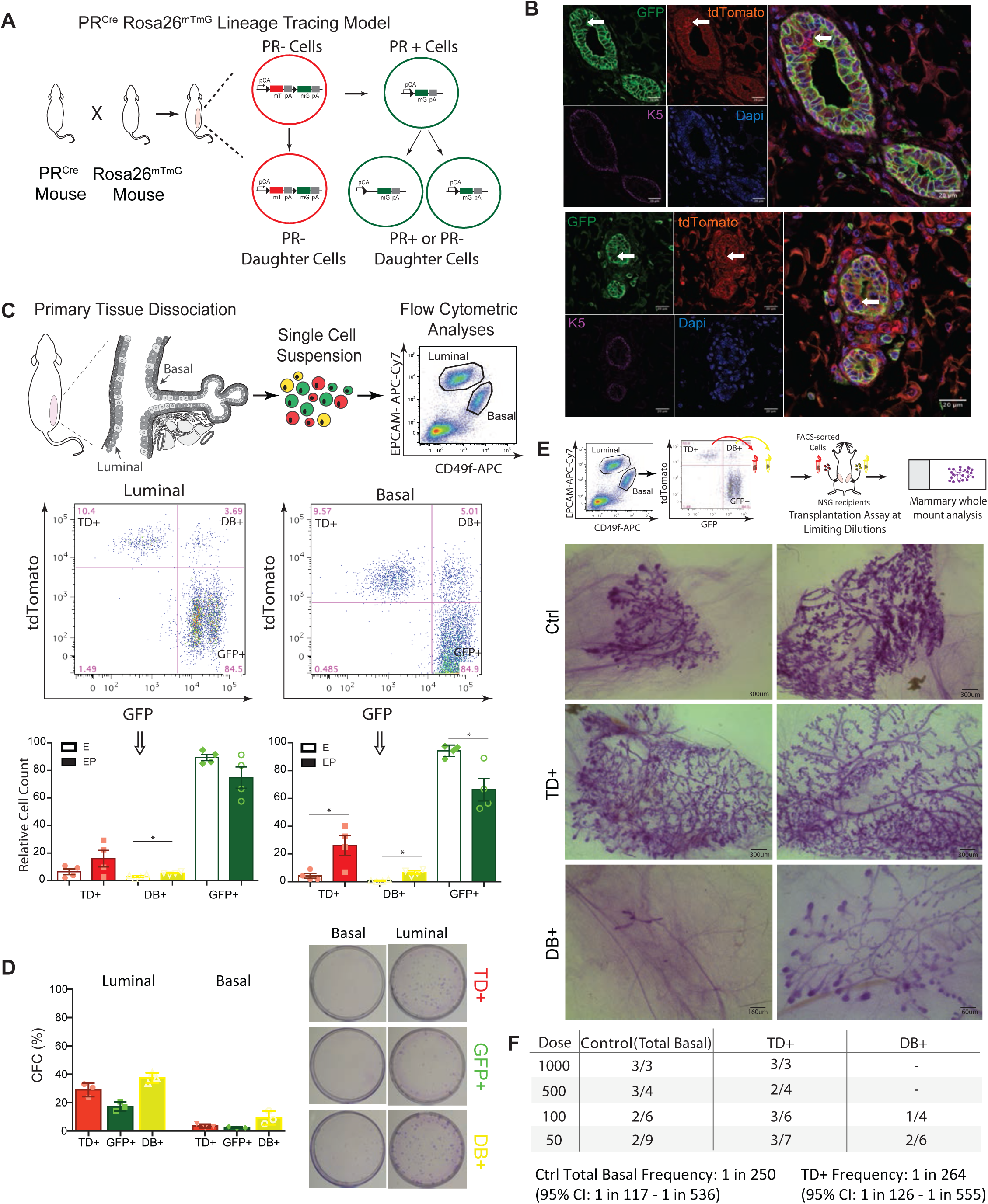
Lineage tracing exposes PR-primed progenitors with repopulation capacity. (A) Schematic of *PR^Cre^ Rosa26^mTmG^* PR-lineage tracing model generated by crossing the *PR^Cre^* GEMM with *Rosa26^mTmG^*reporter GEMM. In the absence of Cre, all cells produce membrane localized tdTomato (TD+) florescence. Upon Cre-mediated excision of tdTomato in PR expressing cells, membrane localized green (GFP+) fluorescence is indelibly expressed in both PR+ cells and all subsequent progenies. (B) Immunofluorescence staining of mammary tissue sections showing the bi-layered mammary duct from adult virgin female *PR^Cre^ Rosa26^mTmG^*mice showing membrane localized tdTomato, PR Cre triggered GFP fluorescence, the K5 basal lineage marker and DAPI as well as merged images. The majority of the epithelium is GFP+ with rare clusters (white arrow top panel) or sporadically scattered (white arrow bottom panel) TD+ cells. (C) Workflow and gating strategy used for flow cytometric analysis of adult virgin glands from the *PR^Cre^ Rosa26^mTmG^*mouse model. Flow plots show a significant proportion of GFP+ epithelium in both the luminal and basal compartments. Quantification of TD+, GFP+ and DB+ cells at baseline following E (hollow bars) and EP treatment (filled bars) in both epithelial compartments. *p<0.05, n=4 per group. (D) Percent of FACS-sorted cells plated that formed colonies from EP treated *PR^Cre^ Rosa26^mTmG^* PR-lineage tracing mice and representative images of corresponding Wright-Giemsa stained CFC plates, n=3 mice. (E) Workflow used for mammary limiting dilution assay of cells from adult virgin glands double FACS-purified from *PR^Cre^ Rosa26^mTmG^*mice. Single cells generated from mammary glands of 10-12-week-old female mice following EP stimulation were injected at limiting dilution into cleared inguinal contra-lateral fat pads of 3-week-old syngeneic recipients and outgrowths scored 5-6 weeks later, at d16-18 of pregnancy. Representative images of wholemount glands obtained after injection of total basal (WT control), TD+ (PR-naïve), and DB+ (td+GFP+ co-expressing) cell subsets. The number of outgrowths reported are pooled from two independent experiments. (F) Limiting dilution assay and quantification of mammary repopulating units based on the number of observed outgrowths out of the total number of glands transplanted with cell subsets.

Bilaterally ovariectomized mice were subjected to classic sex-hormone regimens to mimic the two distinct phases of the reproductive cycle in the mammary gland. Specifically, 17β-oestradiol (E) treatment reflects the baseline gland at the follicular phase of the human reproductive cycle, while 17β-oestradiol+progesterone (EP) phenocopies the proliferative luteal phase and progesterone-triggered expansion of mammary stem/progenitor cells^6^. Flow cytometric analyses confirmed that the majority of cells from both lineages were GFP labelled at baseline (∼91.2% EpCAM^+^CD49^lo^ luminal cells, ∼93.1% EpCAM^+^CD49^hi^ basal cells), with only ∼5.2% of luminal and ∼5.6% of basal cells remaining PR naïve (TD+) (Fig. 1C, S1A). Following progesterone stimulation, this PR naïve cell fraction expanded in both lineages (14.1% luminal, 23.6% basal) (Fig. 1C). These rare PR naïve cells align with the reported ER^−^PR^−^ mammary stem cells situated at the top of the mammary epithelial hierarchy and known to expand upon progesterone exposure^12–14^. We also noted a consistent fraction of PR-primed GFP+/TD+ double positive cells (designated hereafter as DB+) in both lineages, a population typically overlooked in lineage-tracing studies, which expanded in the basal compartment following 17β-oestradiol+progesterone stimulation (Fig. 1C). This unprecedented contribution of PR-primed cells to such a large fraction of both mammary epithelial lineages, especially the basal compartment, challenges our current understanding of mammary biology and alludes to the presence of early bipotent PR-primed progenitors.

### Transplantation of PR-primed DB+ basal subset produces distinct TEB-like outgrowths

We enumerated the clonogenic potential of FACS-purified cell populations (PR naïve TD+, PR lineage-traced GFP+, double positive DB+ cells) from both epithelial lineages in well-established 2-D *in vitro* colony forming assays. Progenitor potential was observed in all three populations, regardless of basal-luminal cell lineage identity or PR status (Fig. 1D). To assess multipotent stem cell capacity, we used the gold-standard limiting dilution assay and transplanted FACS-sorted total basal, TD+ basal or DB+ basal cells in cleared mammary fatpads of pre-pubescent recipients (Fig. 1E, F). The full ductal and alveolar potential of transplanted cells was assessed in mice at gestation d16-18. The PR-naïve TD+ cells consistently produced larger, fuller mammary epithelial outgrowths containing ducts, extensive tertiary branching and robust alveologenesis, consistent with their multipotent capacity at a frequency comparable to the total basal control population. In contrast, the gross size of ductal outgrowths from total basal cells varied, as is usually observed. Most strikingly, DB+ basal cells gave rise to smaller outgrowths, either i) an extremely small duct with limited tertiary branching and alveoli or ii) more extensive ducts with notably engorged ends, similar in morphology to terminal end buds (TEB) seen at puberty. These limited, distinct outgrowths were exclusive to fatpads injected with DB+ cells. This demonstrates the diverse stem/progenitor potential of different cell subsets within the mammary epithelial hierarchy based solely on PR status.

### Basal compartment contains PR+ cells

Next, we directly assessed the status of PR expression in primary mammary epithelium. Generally, ∼1/3 of luminal cells are reported to express ER and/or PR while the basal lineage is considered devoid of hormone receptors. *In situ* immunofluorescence of the PR protein in hormone stimulated mammary tissues was done with validated antibodies that detect both predominant PR isoforms, PRA and PRB. The majority of PR+ cells co-localized with luminal cytokeratin markers (K8/K18) as expected, yet rare PR+ cells co-localized with the basal cytokeratin markers K5 and K14 (Fig. 2A, S2A). Direct intracellular flow cytometry substantiated the PR status of the two epithelial compartments, revealing PR positivity in ∼42% of Lin^−^EpCAM^+^CD49f^lo^ luminal and ∼28% of Lin^−^ EpCAM^+^CD49f^hi^ basal cells (Fig. 2B-D).

**Figure 2.**
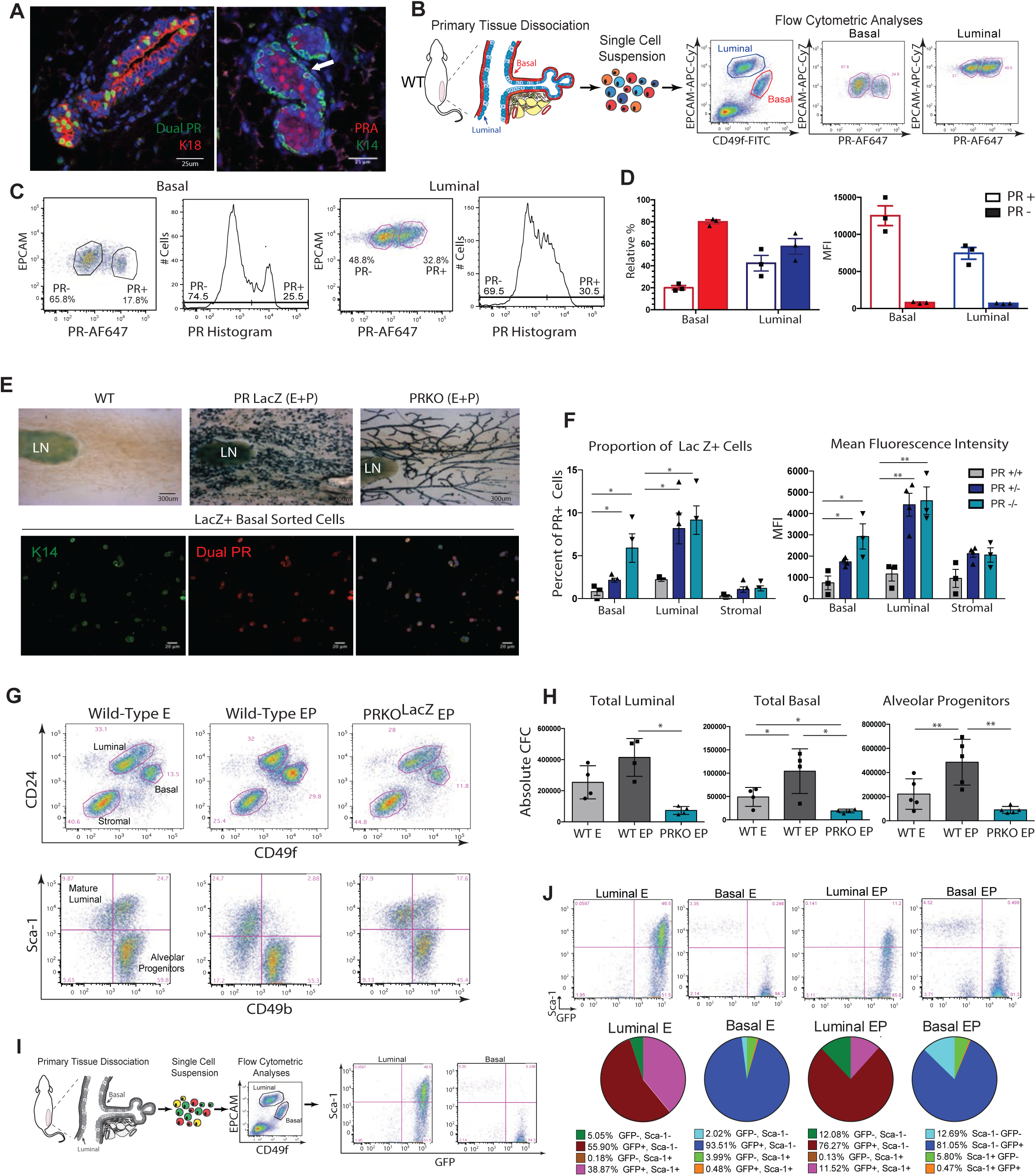
Basal compartment contains PR+ progenitor cells. (A) (Left) Cross-section of bi-layered mammary ducts from WT mice stained with an antibody for total PR (green) and cytokeratin 18 (red), labeling PR+ and luminal cells, respectively. (Right) Cross-section of mammary tissue stained with an anti-PRA antibody (red) and anti-cytokeratin 5 (green, basal marker). White arrow depicts PRA/K5 double positive basal cell (Scale bar = 25 µm). (B) Workflow and gating strategy used for intracellular flow cytometric analysis of PR in wildtype mice. (C) Representative flow plots and histograms showing a clear PR+ subpopulation in both the basal (left) and luminal (right) epithelial subsets. (D) Quantification of PR+ and PR− cells in the luminal and basal epithelial fractions expressed as relative percentage (left) and mean fluorescence intensity of PR staining (right), n=3 mice. (E) (Top) Mammary wholemounts stained with X-gal, a chromogenic substrate of β-galactosidase. Staining of wild-type mammary gland confirming that X-gal staining is specific to tissues with LacZ expression (left). Wholemount from a PR LacZ mouse stimulated with EP (middle) shows strong PR reporter detection in alveoli. In contrast, the PRKO LacZ mouse (right) stimulated with EP lacks alveolar formation. LN= lymph node. (Bottom) Immunofluorescent staining of FACS-purified basal cells based on the PR LacZ signal confirming PR expression at the protein level. (F) Flow cytometry based quantification of proportion (left) and mean fluorescence intensity (right) of PR LacZ+ cells in mammary subpopulations from EP treated PR LacZ littermates WT control (PR+/+), PR LacZ (PR+/−) and PRKO LacZ (PR−/−) mice. *p < 0.05 **p < 0.01, n=3 mice per group. (G) (Top) Flow plots showing the expansion of luminal and basal cells in response to E or EP in WT and PRKO LacZ mice. (Bottom) Flow plots that further segregate the luminal population using the markers Sca-1 and CD49b into distinct progenitor and differentiated cell fractions show a shift in the luminal compartment in response to functional PR signaling. (H) Quantification of the total number of luminal (left), basal (middle) and alveolar progenitors (right) from E and EP treated PR LacZ wild-type littermate control (WT) and PRKO LacZ mice. *p<0.05, **p<0.01, n=4 mice per group. (I) Workflow and gating strategy used for flow cytometric analysis of Sca-1 in *PR^Cre^ Rosa26^mTmG^* mice. Flow plots showing distribution of cells with respect to Sca-1 and the GFP+ PR lineage-trace. (J) (Top) Flow plots showing a strong positive correlation between Sca-1 and the GFP+ PR lineage-trace in the luminal compartment (left) as expected but not the basal compartment (right) at baseline and following EP stimulation. (Bottom) Pie charts emphasizing the expansion in proportions of GFP− cell subsets following EP stimulation compared to E in both the luminal (left) and basal (right) epithelial compartment, n=3 mice per group.

To independently corroborate and quantify PR expression, we leveraged the PR^LacZ^ mouse model. Here, LacZ expression serves as a reporter for both PRA and PRB, enabling quantification of PR expression at the cellular level using LacZ as a surrogate marker. In its homozygous state, it is also a functional PR-knock out (PRKO) GEMM and displays a lack of progesterone-driven alveologenesis and tertiary side-branching^4^, which we confirmed in mammary wholemounts (Fig. 2E). We detected PR^LacZ+^ populations in both epithelial lineages (∼8% luminal, ∼2% basal) and mean fluorescence intensity reiterated PR expression, despite the recognized inefficiency of FDG/LacZ flow cytometry kits (Fig. 2F, S1B). PR^LacZ+^ luminal and basal cells were then FACS-purified and subsequently stained for PR immunofluorescence, verifying FDG/LacZ as a surrogate for PR at the protein level (Fig. 2E, S2B). While a few studies have predicted hormone receptor positivity within the basal lineage, to our knowledge, our multilayered analysis provides the first direct evidence of a PR+ basal subpopulation *in vivo* and *in situ*.

### Basal and luminal progenitor expansions require functional PR

It is known that progesterone stimulates mammary stem and progenitor cell expansion^6,7,25^. Sca-1 and CD49b segregate the luminal population into distinct subpopulations, including a Sca-1^+^CD49b^−^ subset enriched in hormone receptor expressing (HR+) cells with colony forming capacity, an alveolar progenitor-enriched Sca-1^−^CD49b^+^ population expressing milk proteins, and Sca-1^−^CD49b^−^ population only observed during pregnancy^27^. To assess the requirement of functional PRs in mediating progenitor activity, we used the PRKO^LacZ^ model, with its marked abrogation in lobuloalveologenesis. We found progesterone stimulation failed to elicit expansion of total basal or luminal progenitors (Fig. 2F, G). Moreover, CD49b^+^Sca1^−^ alveolar progenitor activity remained low in PRKO epithelium and a shift was noted in the distribution of cells from the Sca-1^+^CD49b^+^ subpopulation to the Sca-1^−^CD49b^−^ luminal subset following sex-hormone exposure (Fig. 2G, H). This is the first report of a PR dependent Sca-1^−^CD49b^−^ subpopulation in the virgin adult gland outside of pregnancy, that is also absent in PRKO^LacZ^ mice.

With our lineage tracing alluding to the presence of an early PR-primed progenitor with demonstrated stem/progenitor activity upon transplantation, we next aimed to ascertain the colony forming capacity of PR+ cells directly (Fig. S2C). Given PR’s intracellular localization, we took advantage of the PR^LacZ^ reporter heterozygous mice, which expresses both functional PR and ß-galactosidase as a surrogate reporter. PR^LacZ^ epithelial subpopulations were FACS-sorted using the FDG-LacZ kits, enabling isolation of PR+ and PR− cells for subsequent live-cell analyses. Specifically, cells from sex-hormone treated PR^LacZ^ mice were FACS-purified into 6 groups for our 2-D colony forming assay: total, PR^LacZ+^, and PR^LacZ-^ populations for both the luminal and basal compartment. As expected, progesterone stimulation increased clonogenicity of total luminal and PR^LacZ-^ luminal populations from baseline levels. Meanwhile, PR^LacZ+^ luminal cell clonogenicity was only observed upon progesterone stimulation. Technical challenges in the viability of basal cells following treatment with the FDG-LacZ kit led to uncharacteristically low progenitor capacity across basal cohorts, regardless of hormone treatments (Fig. S2C). Nevertheless, flow plots showed that PR+ basal cells were enriched within the most CD24^+^CD49f^+^ tip of the basal population, a subset reported to be enriched in mammary repopulating units (MRU^12^, Fig. S2D). FACS-purification and subsequent qRT-PCR analysis of the MRU-enriched basal fraction compared to the rest of the basal population revealed higher PR expression and Cyclin D1 expression, a cell cycle protein required for progesterone-induced proliferation in a cell-autonomous manner (Fig. S2E). This provides direct demonstration of a PR+ population with clonogenic capacity, underscoring a unique role for PR in sex-hormone driven epithelial cell dynamics.

Finally, Sca-1 has also been shown to closely correlate with ER expression, often serving as a surrogate for overall hormone receptor positivity in mammary epithelial cells^27^. In our *PR^Cre^ Rosa26^mTmG^* lineage tracing model, we found a strong correlation between Sca-1+ and GFP+ luminal cells, with back-gating showing a distinct, hormone-responsive luminal subset (Fig. 2I, J). In contrast, GFP+ basal cells did not correlate with Sca-1 expression. Thus, basal and luminal GFP+ populations are distinct beyond their lineage identity, despite both being hormone responsive cells.

### Proteomes confirm PR+ basal cells are a discrete subset within basal lineage

Next, we profiled the mammary epithelial subpopulations using low-input proteomics^28^ on equivalent numbers of matched, FACS-purified PR+ and PR− cells from basal and luminal lineages from sex-hormone treated wild-type mice (Fig. 3A). Of the 4,452 detected proteins, 2,857 were consistently detected (≥4 of 6 mice) in all four cell types; 524 and 160 proteins were unique to luminal and basal cells, respectively (Fig. 3B). Principal component analysis indicated mammary cell lineage as the dominant clustering feature, with secondary segregation based on PR status (Fig. S3A). Lineage markers (basal: ITGB1, ITGA6, VIM, KRT5, KRT14; luminal: KRT8, KRT18, KRT19, EPCAM, GATA3, ITGA2, MFGE8) recapitulated known expression patterns (Fig. 3C, D) and were among the most differentially expressed proteins between basal and luminal lineages in PR+ and PR− subsets. Many differential proteins were driven by lineage identity (Fig. S3B), consistent with the divergent nature of luminal-basal cells. 98% of the 1,303 significantly different proteins between basal and luminal cells had the same directionality in both PR+ and PR− subsets (Fig. 3E). Of the 4 cell types, the basal PR+ and basal PR− cells are the most similar (r =0.97), followed by luminal PR+ and luminal PR− cells (r =0.96) (Fig. S3C). The correlation between cells of luminal and basal lineages were more dissimilar independent of PR status (r of 0.80-0.82) (Fig. S3C). These proteomic analyses demonstrate that basal PR+ cells are *bona fide* cells of the basal lineage and not an aberrant subset of the luminal lineage.

**Figure 3.**
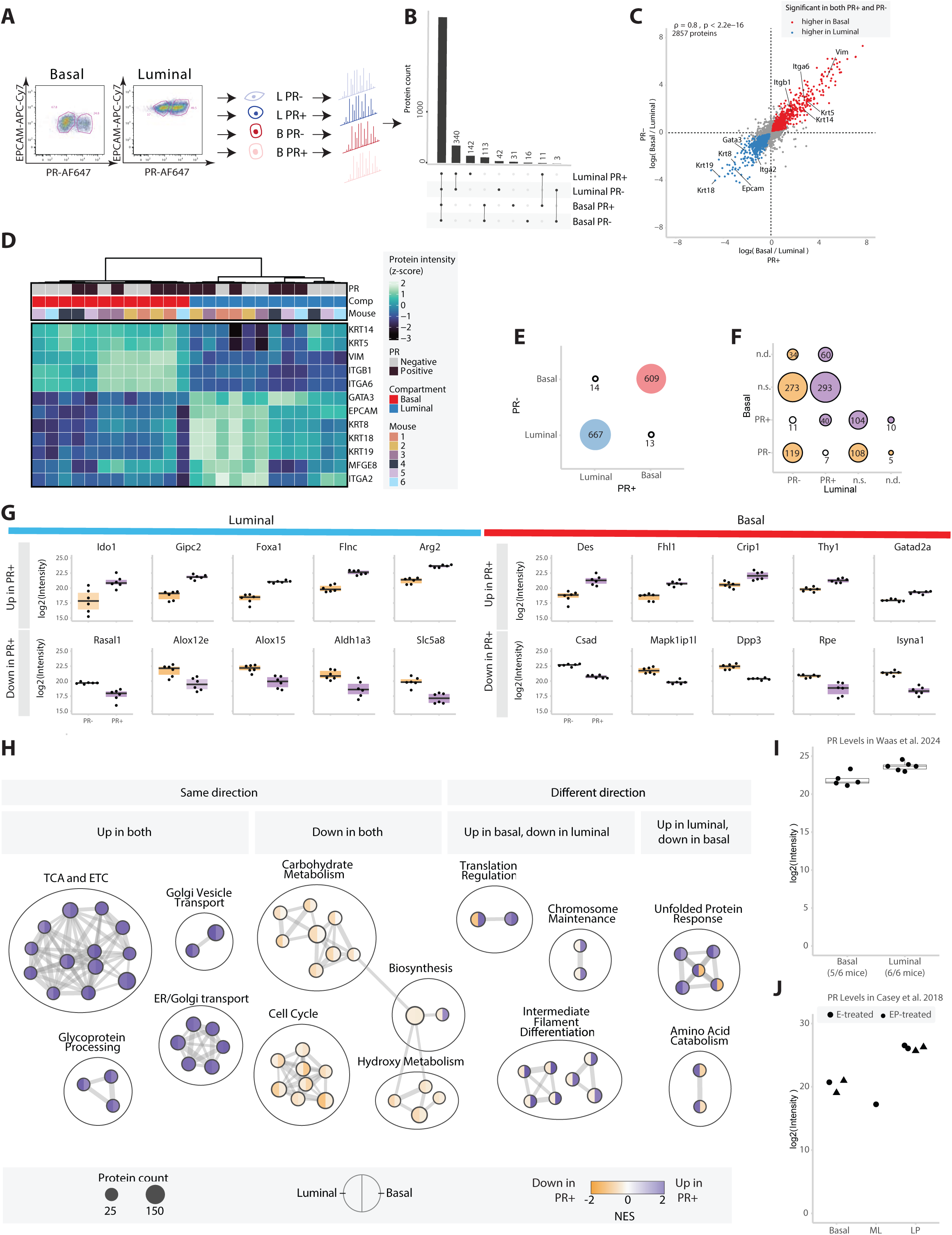
Proteomes confirm PR+ basal cells are a discrete subset within basal lineage. (A) Low input proteomics workflow used to profile PR+ and PR− cells of luminal and basal lineage using DROPPS, n = 6 mice. (B) An UpSet depicting the overlap in the proteins consistently detected (≥4 of 6 mice) in the different sample types. (C) Protein expression fold changes between basal and luminal cells from PR− cells plotted against those from PR+ cells. Known basal and luminal lineage markers are labeled. Color signifies proteins which are significantly different basal and luminal for both the PR− and PR+ subsets. The Spearman’s correlation of the fold changes is shown. (D) Heatmap of z-scaled protein log_2_(Intensity) of known basal and luminal markers. (E,F) Bubble plot showing the number and directionality of proteins significantly different between (E) basal and luminal lineages in PR+ and PR− subsets and (F) between PR+ and PR− subsets in basal and luminal lineages; n.s. = not significant, n.d. = not detected. (G) The top 5 differentially expressed proteins between PR+ and PR− subsets in each direction within basal and luminal lineages. Boxplots show median and interquartile range. (H) Cytoscape enrichment network showing pathways which are different based on PR expression in the basal and luminal lineages. (I, J) Proteomic data from (I) Waas et. al (n = 6 mice) and (J) Casey et al. (each point represents pool from 3 mice) depicting the detection of PR in mouse basal cells. Boxplots show median, interquartile range, and 95% confidence interval estimate.

In contrast, protein expression differences associated with PR are divergent between lineages (r =0.36, Fig. S3D). Of the 393 proteins higher in luminal PR+ cells vs PR− cells, 15% and 75% were not detected or were not significantly different, respectively in basal cells (Fig. 3F). Only 40 proteins were higher in both PR+ subsets compared to the respective PR− subsets (Fig. 3F). The top differentially expressed proteins across PR subsets were distinct between basal and luminal cells (Fig. 3G). Despite these differences, there are many pathways significantly associated with PR in both luminal and basal lineages (Fig. 3H). The most prominent similarity between lineages was enrichment for *TCA and ETC* in PR+ cells in contrast to enrichment for *carbohydrate metabolism* in PR− cells (Fig. 3H). This hints at the hormonal influence on breast mitochondrial biology and metabolism, aligning with human mammary proteomes that showed similar sets of differential pathways in pre- versus post-menopausal women^29^. Some pathways were enriched in opposing directions based on PR status, for example *intermediate filament differentiation* was enriched in basal PR+ and luminal PR− compared to basal PR− and luminal PR+, respectively (Fig. 3H). This suggests PR expression may be associated with different functional requirements between luminal and basal lineages. Pathways associated with basal and luminal lineage identity are essentially identical across PR subsets (Fig. S3E, F). Terms including *fatty acid metabolism* and *translation initiation* were enriched in luminal cells and terms for *cell substrate adhesion* and *muscle contraction* were enriched in basal cells in both PR+ and PR− subsets (Fig. S3E). Altogether, these analyses indicate that lineage differences are largely conserved independent of PR status. However, the proteomic landscape that separates a luminal PR+ from a PR− cell is distinct from what renders a basal PR+ from a PR− cell. This points to PR playing different roles in basal and luminal cells. Finally, we interrogated two published mammary epithelial proteomic datasets for additional evidence of PR in basal cells. In Waas et al.^28^, PR was detected in the basal lineage in 5 of 6 mice (Fig. 3I) and in Casey et al.^30^, PR was detected in 3 of 4 pools of cells from the basal lineage (Fig. 3J). Altogether these observations demonstrate that PR has been detected at the protein level in basal cells across multiple independent studies.

### Analysis of the human breast corroborates PR+ basal cells

We examined published datasets from global OMICs efforts looking for PR in the basal compartment of the murine mammary gland and human breast. To assess the frequency and characteristics of the PR+ basal population, we interrogated sc-RNAseq studies and found only 74 luminal PR+ cells and 9 PR+ basal murine cells in Pal et al. (2017)^31^; 921/10015 PR+ basal and 4246/13169 PR+ luminal murine cells in Bach et al (2017)^32^; and a total of 59/3252 PR+ basal and 2281/29694 PR+ luminal human cells in Gray et al (2022)^33^ (data not shown). Although fewer than expected PR+ luminal cells were observed, reflecting a gross underrepresentation of HR+ cells, PR+ basal cells were readily identified in all datasets. We then leveraged the recent Human Breast Cell Atlas epithelial dataset (Kumar et al. 2023)^34^ where 113/126 breast samples contained PR+ basal cells and ∼7% of the basal population expressed *PGR* (Fig. 4A, S4A). Comparatively, 123/126 breast samples contained PR+ mature luminal (‘lumhr’) cells and ∼26% of the lumhr population expressed *PGR* (Fig. S4B, C). Pathway analysis (Fig. S4D) of PR+ versus PR− basal cells showed Reactome Signaling by Wnt as the most enriched term (Fig. 4B), with the inhibitor of WNT-dependent transcription *SFRP1* and the putative mammary progenitor cell marker *SOX9* as the top Wnt related genes driving this enrichment (Fig. 4C).

**Figure 4.**
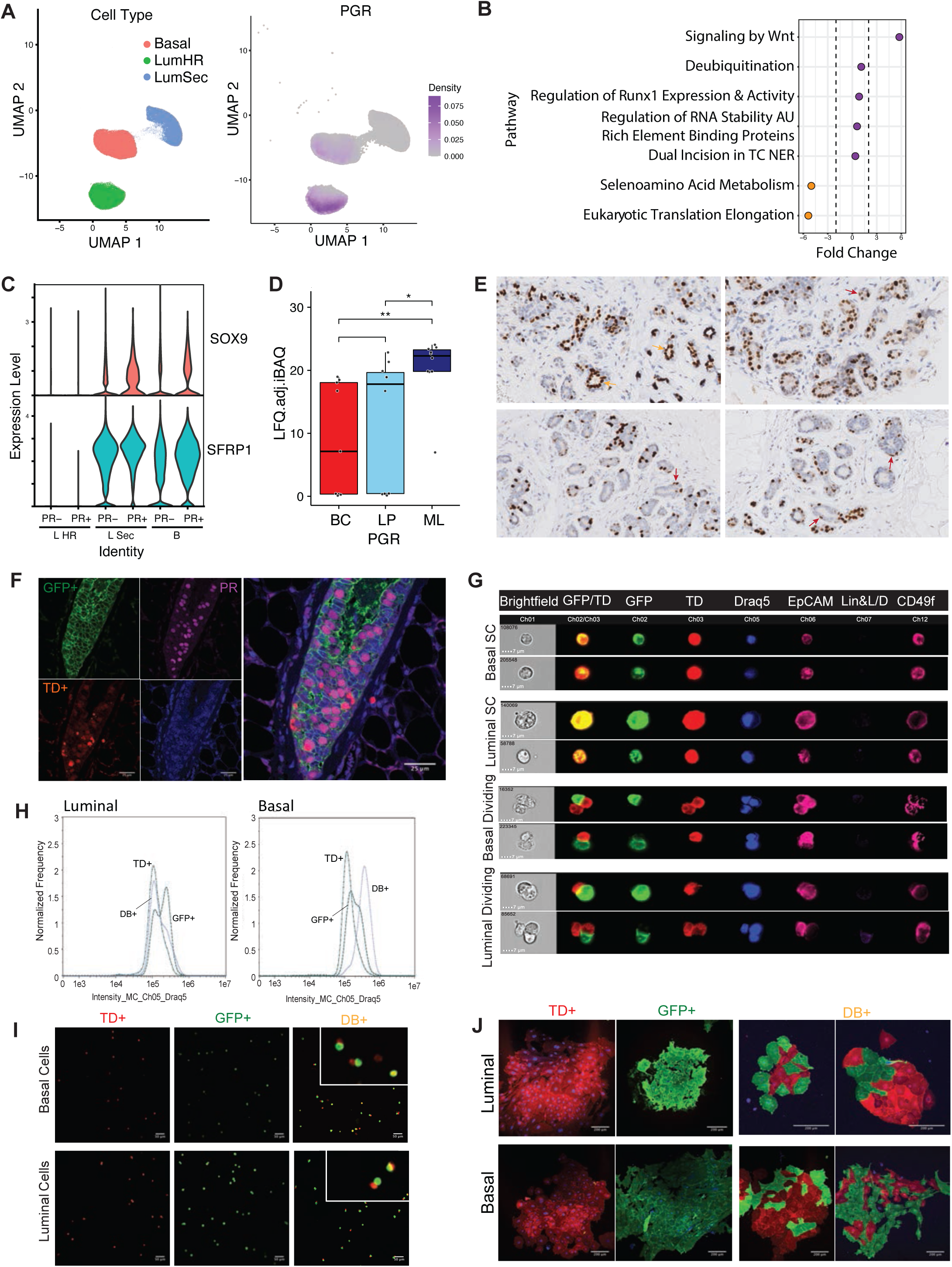
Human PR+ Basal Cells and Asymmetric Division. (A) Uniform manifold approximation and projection (UMAP) of epithelial subset from Kumar et al (2023) showing cell clusters coloured by cell type (left) and gene expression intensity of PGR (right), including in the basal cell cluster. (B) Pathway analysis of PGR+ basal cells compared to PGR− basal cells from Kumar et al (2023) showed Reactome Signaling by Wnt as the most enriched pathway in PGR+ basal cells. (C) Violin plot showing expression of the top two Wnt related genes from the pathway analysis in mammary epithelial subclusters based on PGR status in Kumar et al (2023). These included the inhibitor of WNT-dependent transcription SFRP1 (bottom) and the putative mammary progenitor cell marker SOX9 (top). (D) Box plot showing PR peptide detection in FACS sorted human breast epithelial subpopulations (BC, basal cell; LP, luminal progenitor; ML mature luminal cell). Proteomes included samples from both pre-and post-menopausal women. (*p=0.05, **p=0.01). (E) PR immunohistochemical staining of human breast tissues. Lumen adjacent PR positive cells are widely detected as expected (orange arrow) but a number of basally positioned PR positive cells are also observed (red arrows). (F) Cross-sections of the bi-layered mammary duct from adult virgin female *PR^Cre^ Rosa26^mTmG^* PR− lineage tracing mice depicting the expression of membrane localized tdTomato (TD+), PR^Cre^ induced GFP fluorescence, PR and DAPI (blue), as well as the combined stains. (G) A panel of representative Amnis ImageStream®X images displaying GFP and tdTomato expression in cells from the mouse luminal and basal compartments as marked by the set of established cell surface markers EpCAM and CD49f. Two subsets of cells were detected: i) a subset of single cells which are double positive cells and express both GFP and TD signals (top 4 rows) and ii) a distinct subset of cells that appear attached/recently divided with a single GFP+ PR-primed cell alongside a TD+ PR-naïve cell (bottom 4 rows). (H) Histogram depicting the distribution of the DNA content using DRAQ5 in TD+, GFP+ and DB+ double positive cells by Amnis imaging flow cytometry, n= 3 mice. (I) Confocal immunofluorescent images following cytospin analysis of FACS-purified TD+, GFP+, and DB+ cells from the *PR^Cre^ Rosa26^mTmG^* PR-lineage tracing mice. Two subsets of cells were detected: i) a subset of unique single cells that were double positive for both eGFP+ and tdTomato+ (appearing yellow, DB+) and ii) a subset of paired cells, with one TD+ PR-naïve cell immediately adjacent to and/or attached to a GFP+ PR traced cell. (Scale bar = 20um). (J) Fluorescent imaging for GFP and tdTomato in colonies from FACS-sorted TD+, GFP+ PR-traced, and DB+ cells from the luminal and basal compartments following EP stimulation and plated in standard CFC assays, n= 3 mice.

At the protein level in the human breast, we observed PGR peptides in 5/9 patients in our published basal cell proteomes, which included both pre- and post-menopausal individuals (Fig. 4D). Finally, we sought to discern basal/myoepithelial PR+ cells in situ in primary breast tissue sections. Vacuum Assisted Biopsy derived breast biopsies from a total of 17 premenopausal participants in the Breast Cancer Anti-progestin prevention study-1 (NCT02408770) were stained with a robust, clinical grade PR antibody for immunohistochemistry. We observed a subset of non-luminal PR+ cells based off cell morphology and cell orientation across multiple patient samples, despite intra-patient and inter-patient heterogeneity in PR expression (Fig. 4E). These basally positioned PR+ cells appeared more rounded (basal-like) or more elongated (myoepithelial-like), instead of columnar (luminal-like), and were often situated parallel to the ductal lumen, similar to myoepithelial cells, rather than perpendicular (Fig. 4E). These analyses of primary human breast epithelium reiterate the presence of PR+ basal cells.

### PR-primed progenitors give rise to PR+ and PR− progeny and divide asymmetrically

Since PR-primed cells are labelled indelibly by GFP in *PR^Cre^ Rosa26^mTmG^* mice, tissues were immunofluorescently co-stained with a PR antibody to determine the hormone receptor protein status of PR-primed cells as well as all progeny. Notably, not all GFP+ cells co-expressed PR, indicating that not all PR-primed cells or their progeny continue to express PR (Fig. 4F). In contrast to previous reports that all ER-lineage traced cells and progeny maintain ER expression, our unique observations deviate from current models of hormone receptor patterning in mammary epithelium, not only based on lineage-identity but also the progenitor-progeny hierarchy.

As noted in our PR-lineage tracing experiments above, we consistently observed a dynamic population of DB+ cells in both luminal and basal compartments which expanded upon sex-hormone treatment. Moreover, DB+ cells possessed progenitor potential in vitro and the ability to form mammary outgrowths upon transplantation (Fig. 1D, E) and therefore sought to analyze them further. Complementing traditional flow cytometry, high throughput imaging flow cytometry confirmed this populations of cells in both lineages to indeed be single cells based on cell size and shape that expressed both GFP and tdTomato, rather than cell doublets (Fig. 4G). Notably, a subset of this population in both lineages appeared to be either dividing or have recently divided, based on DNA content and cell morphology (Fig. 4H, I). FACS-purification of this DB+ population from each lineage followed by cytospin analysis confirmed these to be distinctive, i.e. consisting of a subset of single cells co-expressing GFP and tdTomato as well as a subset of recently divided cells that asymmetrically produced one TD+ and one GFP+ cell (Fig. 4I). Finally, we compared clonogenic potential of FACS-purified cell populations (PR-naïve TD+, PR-traced GFP+, and GFP+/tdTomato DB+ cells in each lineage) using 2-D colony forming assays. TD+ cells largely gave rise to TD+ colonies while GFP+ traced cells formed colonies exclusively marked with the GFP trace, as expected. Most strikingly, DB+ cells exhibited GFP+ colonies (consisting entirely of GFP+ cells) or one-of-a-kind mixed colonies, which contained integrated GFP+ and TD+ cells, likely arising from asymmetric cell division (Fig. 4J). These data show that distinct progenitor populations exist based on PR status in the mammary epithelial hierarchy in both cell lineages.

## DISCUSSION

In a constitutive PR lineage-tracing mouse model, we reveal the unexpected contribution to all the mammary epithelial compartments, alluding to the presence of PR-primed progenitors. We isolate PR-primed progenitors from both epithelial lineages and demonstrate their unique stem potential. We identify a discrete de novo PR expressing population in the basal compartment and dissect its proteomic landscape. These data redefine our understanding of the mammary epithelial hierarchy and opens possibility for direct autocrine hormonal responses in both mammary lineages.

The discovery of single self-renewing mammary stem cells that functionally generate the full ductal tree and give rise to all epithelial cell types (luminal and basal, HR+ and HR-), illuminated the field to the immense regenerative potential inherent to the breast,^84,85^. Luminal and basal breast epithelium in the same region possess identical chromosomal alterations, implicating shared ancestry^35^ and scRNA-seq of the human breast found a continuous lineage hierarchy, connecting the basal with the two luminal branches^36^. Furthermore, select lineage tracing reports document the presence of a bipotent basal population^37,38^. However, other lineage tracing studies have challenged this classical view, revealing combined in situ action of unipotent basal and luminal progenitors in maintaining the mammary gland^39–41^. Lineage-tracing of ER expressing cells indicated luminal ER+ stem/progenitors exclusively sustain all ER+ mammary epithelial cells^26,41^. Meanwhile, PR transcripts have been found in the bipotent enriched fraction of normal human breast cells and EpCAM^+^CD49f^+^ fraction of normal murine mammary cells^6,42,43^. Early reports of label-retaining mammary epithelial cells (LRECs), that keep their template DNA strands during mitosis, showed PR expression in up to ∼30-40% of cells^44,45^ and a large fraction of LRECs from the normal breast transplanted into mice likewise expressed PR^46^. CD10^+^ bipotent progenitors similarly contain PR+ cells ^9,47–49^. PR immunofluorescence was noted in a fraction of SMA^+^K14^+^p63^+^ human basal cells and in basally positioned K8+ progenitor cells, where the PRB isoform was implicated in contrast to the more readily detected and studied PRA isoform^13^. Our study provides credence to these often overlooked studies, the classical notion of an early bipotent mammary epithelial stem/progenitor, and highlights a far more complex role of PR expressing cells in the mammary epithelial hierarchy than previously anticipated.

Knowledge of parent-progeny cell relationships in the mammary epithelial hierarchy spawned the cancer cell-of-origin concept and has proven imperative for enabling breast cancer interception strategies. The discovery that progesterone induced stem cell expansion^6,7^ through paracrine mechanisms^22,24,50^, created a paradigm shift in our understanding of increased breast cancer risk associated with increased sex-hormone exposure ^41,51–53^. Specifically, targeting essential paracrine effectors (anti-RANKL, NCT04711109, NCT04067726) and progesterone itself (BCAPPS-1 NCT02408770) have gained traction as a means to limit aberrant epithelial proliferation^54^ and mitigate breast cancer risk. Our discovery of a PR+ basal population opens new avenues, presenting a novel basal target population of interest. It raises possibilities of a direct autocrine basal cell response to progesterone that can bypass luminal paracrine effectors, rendering the direct inhibition of PR a more attractive approach. Overall our study provides novel insights into PR regulation of the normal breast hierarchy that should hasten the development of effective breast cancer preventive strategies.

## METHODS

### Mouse Models and Hormone Treatments

PR^Cre^ (Stock# 017915) and Rosa26^mTmG^ (Stock# 007676), mice were obtained from the Jackson Laboratory. PR Lac Z reporter mice were generously donated from Dr. John Lydon’s Lab at the Baylor College of Medicine in Houston, Texas. C57BL/6 and FVB 8-12-week-old virgin female mice were purchased from Charles River Laboratories as needed. For transplantation experiments, NSG female mice were cleared of endogenous mammary outgrowths at 20/21 day-old. For hormone controlled experiments, virgin female mice of approximately 10 weeks of age were ovariectomized bilaterally. Following a one-week recovery period, mice were subjected to a classic sex-hormone regimens with either a 14-day slow-release pellet containing 0.14 mg 17-β estradiol (E) or a 0.14 mg 17β-estradiol plus 14 mg progesterone (EP; Innovative Research of America). Upon the completion of a treatment, mice were sacrificed and three pairs of mammary glands (second, thoracic, and fourth inguinal) were collected for subsequent analyses. All mice were housed in a standard, controlled environment where the light cycle in the rooms was set to 12 h on and 12 h off with a 30-min transition, the temperature was 21–23 °C, and humidity at 30– 60%. NC3Rs ARRIVE, Canadian Council for Animal Care guidelines and protocols approved by the Animal Care Committee of the Princess Margaret Cancer Centre were strictly followed.

### Mammary Cell Preparation

Single-cell suspensions were prepared by mincing freshly harvested mammary glands and incubating them in DMEM/F12 media with 750 U/ml collagenase and 250 U/ml hyaluronidase (Stem Cell Technologies, 07912) at 37°C for 1.5 hours.^12^ Samples were vortexed at the 1 and 1.5 h mark. This was followed by treatments with ammonium chloride (Stem Cell Technologies, 07850), 0.25% trypsin-EDTA (0.25%, Stem Cell Technologies, 07901) and 5 U/mL dispase (Stem Cell Technologies, 07913) plus 50 ug/ml DNase I (Sigma, D4513) in Hank’s Balanced Salt Solution (HBSS; Gibco, 14025092) supplemented with 2% FBS (Gibco, 12483020). The resulting single cells were filtered through a 40 μm cell strainer (Fisher Scientific, 22-363-547). The cells were washed between each of the above steps with 10 ml of Hank’s Balanced Salt Solution with 2% FBS, and centrifugation at 1200 rpm.

### Fluorescence-activated cell sorting (FACS)

Common to all staining plans, cells were stained using the murine chimera biotinylated StemSep cocktail (StemCell Technologies, 19849) along with an anti-CD31 antibody to exclude CD45+/Ter119+ hematopoietic and CD31+ endothelial cells upon secondary conjugation with streptavidin-PE-Cy7. Alternatively, directly conjugated CD45, Ter119 and CD31 antibodies were utilized to exclude these cells. Propidium iodide or DAPI were used as viability dye (Sigma, D9542). To analyze PR Lac Z signal, the FluoReporter Lac Z Flow Cytometry Kit (Fisher Scientific, F1930) was used, which utilizes the fluorescent β - galactosidase substrate fluorescein di-V-galactoside (FDG). For this staining protocol, cells were initially stained using an anti-CD49f-APC (clone GoH3, BD pharmigen) and CD24-PE (clone M1/69, eBiosciences) to resolve the mammary epithelial compartments, then incubated for 10 min at 37°C with chloroquinone to prevent lysosomal β-galactosidase degradation of FDG. FDG is then added to cells and continued to incubate at 37°C for an additional minute before it is placed on ice. Ice cold staining medium (4% FBS, 10 mM HEPES in PBS) is then added to each sample. Following centrifugation at 1200rpm, the cells are resuspended in HANKS BSS containing propidium iodide. Cells were sorted into HANKS BSS or PBS for subsequent analyses as needed using the BD FACSAria Fusion.

### Flow cytometry

Mammary epithelial subpopulations, were stained with a specific cocktail of antibodies. Dead cells were excluded following doublet exclusion using DAPI or Zombie UV Fixable Viability Kit (BioLegend, 423107) according to manufacturer’s instructions. Antibodies specific for TER119 (PECy7 or eFluor450), CD31 (PECy7 or eFluor450), CD45 (PECy7 or eFluor450), EpCAM (APCCy7 or PE), CD24(PE), CD49f (FITC or PECy7), CD49b (PE) and Sca-1 (APC or Brilliant Violet 711) were used. Lineage (Lin) positive cells were defined as Ter119^+^CD31^+^CD45^+^. Mouse mammary cell subpopulations were defined as: total basal (Lin^−^EpCAM^lo-med^CD49f^hi^ or Lin^−^CD24^lo-med^CD49f^hi^) or total luminal (Lin^−^EpCAM^hi^CD49f^lo^ or Lin^−^CD24^hi^CD49f^lo^). For direct intracellular flow cytometry, ∼500ul of 4% PFA was added to the cell suspension and incubated for 10 min at room temperature (RT) then washed with either Hank’s buffer or PBS before incubation with 0.1% Triton-X for 5min at RT. After wash the PR antibody (clone F2, Santa Cruz, 1:200) was added for 30 min on ice. Following a final wash cells were resuspended in PBS with or without fixable live/dead zombie UV dyes to distinguish live/dead cell. Flow cytometry was performed on BD Fortessa Analyzer or from FACS on the BD FACS Aria Fusion and results analyzed with FlowJo software (Tree Star, Inc).

### Mammary wholemount analysis and X-Gal staining

The fourth inguinal mammary glands were removed and carefully spread out on a microscope slide. Slides were then fixed in Carnoy’s solution overnight (ON) at RT. The glands were then rehydrated through a graded EtOH gradient series of 70%, 50%, 30% and 10% x2 for 10 min each, and two 5 minute washes in distilled water. Slides were then immersed in Carmine Alum stain ON at RT. Finally, slides were dehydrated with serial EtOH washes of 75%, 95%, and 100% for 15 min each, and a 5 min xylene wash before mounting with Permount and a glass cover slip. For mammary x-gal wholemounts, the 4^th^ inguinal glands were fixed in 2% paraformaldehyde (PFA), pH7.4, 2h at 4°C, washed in PBS for 30 min x3, and immersed in X-Gal staining solution (chromogenic β-galactosidase substrate 5-bromo-4-chloro-3-indolyl-beta-D-galacto-pyranoside, 1.3 mM MgCl_2_, 15 mM NaCl, 5 mM K_3_Fe(CN)_6_ 3H_2_0, 5 mM K_4_Fe(CN)_6_ 3H_2_0, 0.02% NP-40, 44 mM HEPES; pH7.9) at 0.05% wt/vol X-gal) at 37°C for 14-16 h in the dark. After three 30 min washes in PBS, glands were immersed in an EtOH series of 70%, 95% and 100% for 15 minutes each. Finally, the glands were incubated in xylene ON and mounted with Permount.

### Histology and Immunofluorescence

Freshly harvested mouse mammary glands were fixed in 4% PFA at 4°C ON and subsequently stored in 70% EtOH. Toronto Center for Phenogenomics performed paraffin-embedding and tissue sectioning services. Tissue-section slides underwent deparaffinization and rehydration prior to antigen retrieval in Decloaking ChamberTM Pro for 30 minutes at 121°C in Reveal Decloaker (Biocare Medical pH 6.0 citrate buffer). The sections were incubated with blocking buffer (20% goat serum, 4% BSA, and 0.3% Triton-X in PBS) for 1h at RT. The sections were stained with a cocktail of primary antibodies that were diluted in the same blocking buffer ON at 4°C. Next day, the tissue sections were washed in 0.1% Tween-20/ PBS and then incubated with secondary antibodies diluted in blocking buffer for 1h at RT. The sections were mounted in Anti-fade ProLongTM Gold with DAPI (Life Technologies, P36935). Primary antibodies used include a mouse anti-keratin 14 (cat. sc-53253, Santa Cruz), a mouse anti-keratin 18 (cat. 10R-C161a, Fitzgerald), a rabbit anti-keratin 5 (Biolegend, Poly19055), a rabbit anti-keratin 8 (Biolegend, Poly19056), and a dual rabbit anti-PR (sc-7208, Santa Cruz), a chicken anti-beta galactosidase (cat. ab9361, Abcam), a mouse anti-PRA (clone hPRa7, cat. MA5-12658, ThermoFisher), a mouse anti-PRB (clone hPRa6, cat. MA5-12653, ThermoFisher) and anti-PR dual isoform antibody (sc-166170, clone F-2, Santa Cruz). Secondary antibodies include anti-mouse conjugated to AlexaFluor 647 (Cedarlane, 115-605-166, 1:300), anti-chicken-Cy3 (Cedarlane, 703-165-155, 1:500), and anti-rabbit-AlexaFluor 488 (ThermoFisher, A-11008). All tissue immunofluorescence images were acquired on a Zeiss LSM 700 confocal microscope using a 40X oil-immersion objective lens. At least three or four Z-planes were acquired per field of view and final images generated using ImageJ software.

### Mouse CFC assay

FACS-purified populations (350 cells) were seeded together with 20,000 irradiated NIH 3T3 cells in 6-well plates or 6 cm dishes. Cells were cultured for 7 days at 5% oxygen in EpiCult-B mouse medium (Stem Cell Technologies, 05610) supplemented with 5% FBS, 10 ng/ml EGF, 20 ng/ml basic FGF, 4 μg/ml heparin, and 5 μM ROCK inhibitor (Millipore). On day 7, plates were subsequently fixed in a 1:1 acetone:methanol (v/v) for 1 min and stained using Wright’s Giemsa stain (Fisher Diagnostics, 264-983). Total number of colonies per plate were scored manually using a Leitz dissecting microscope.

### In vivo transplantation assay

Double FACS-purified total basal, TD+ basal cells or DB+ (double positive) basal cells were resuspended in a 10 μL volume containing 50% Matrigel and 50% Hanks’ balanced salt solution plus 2% FBS and 0.02% Trypan blue (Sigma). Cells (10,000, 5,000, 1,000 and 500) were injected into cleared 4^th^ inguinal mammary fat pads of pre-pubescent, 21-day-old NSG female recipients. Specifically, control cells and a cell subset of interest were transplanted into the two contra-lateral 4^th^ inguinal glands of the same recipient mouse to preclude a differential effect on take rate arising from variation in the hormonal milieu of the host recipient. After 5 weeks, mice were mated and mammary glands were dissected on gestational d16-18 to assess the ductal and alveolar potential of the transplanted cells. Carmine alum-staining was done for wholemount analysis. Fat pads were scored as positive or negative depending on the presence or absence of ductal and alveolar outgrowth. The frequency of MRUs in single cell suspensions from the specific injection groups was calculated using Poisson statistics and the method of maximum likelihood using L-Calc software (Stem Cell Technologies).

### RNA isolation and real-time PCR analysis

RNA was isolated from FACS-sorted mammary epithelial cells using the Arcturus PicoPure RNA Isolation Kits following the manufacturer’s protocol (Applied Biosystems). First strand cDNA was reverse transcribed from the isolated RNA using the SMARTer PCR cDNA Synthesis Kit (Clontech). For epithelial subpopulations with a limited number of cells resulting in low levels of RNA, the first strand was amplified using the Advantage2 PCR kit (Clontech) and the optimal number of amplification cycles for each sample was individually determined by analyzing amplified cDNA aliquots on an ethidium bromide-stained 1.2% agarose gel. Relative quantification of Real-Time PCR (Ct) was performed on 4 ng of cDNA using an ABI PRISM 7900HT Sequence Detection System and Taqman gene expression assay mixes containing unlabelled PCR primers and FAM labeled Taqman MBG probes (Table 1-1). Expression levels of target genes were normalized to *β-actin* transcripts. Previously published, custom primer probe sets were used to detect mouse PRB and PRAB transcripts.^6^

### AMNIS Acquisition and analysis

Freshly dissociated mouse single cells were stained with a fixable viability Zombie UV Dye (BioLegend, 423107, 1:100) and a cocktail of cell surface markers including: biotin-conjugated CD45 (eBioscience, 13-0451-82, 1:800), CD31 (eBioscience, 13-0311-8, 1:200), Ter119 (eBioscience, 13-5921-81, 1:100) which were subsequently labelled with secondary conjugate streptavidin-eFluor 450™ (eBioscience, 48-4317-82, 1:500); anti-EpCAM-APC-eFluor® 780 (eBioscience, 47-5791-82, 1:200), anti-CD49f-PE/Cy7 (BioLegend, 313622, 1:100), and endogenous cell fluorescence in eGFP and tdTomato. After washing, cell surface stained cells were fixed in 4% PFA for 10 min at RT, washed in PBS, and stored ON at 4°C. The next day, the cells were permeabilized with 0.1% Triton-X/PBS for 5 min at RT, washed and nuclear DNA was stained by DRAQ5™ (Thermo Fisher Scientific 62251, 2.5 μM). Imaging flow cytometry was performed on an ImageStream®X Mark II (MilliporeSigma) with INSPIRE® software. Stained cells were resuspended in a volume of 50 μl PBS + 1% FBS in a 1.5 mL low retention microfuge tube (Sigma, T4816). Samples were then acquired on a 5 laser 12 channel ImageStream®X Mark II imaging flow cytometer at 60X magnification following ASSIST calibration (Amnis® Corporation). Channels 1, 2, 3, 4, 6, 7, 9, 11 and 12 were used for acquisition along with lasers 405 nm (100 mW), 488 nm (150 mW), 561 nm (200 mW), 592 nm (300 mW) and 642 nm (150 mW) for excitation. A bright-field (BF) area lower limit of 50 μm^2^ was used to eliminate debris and calibration beads during sample acquisition, while samples were collected in a series of 50×103 event raw image files (.rif). For single stained compensation controls, BF illumination was turned off and approximately 3000 events within the positive signal fraction were acquired. An initial compensation matrix was generated by loading the single stained raw image files into the IDEAS compensation wizard (IDEAS® version 6.2) with further refinements to the compensation matrix made as necessary through manual adjustment. Once generated, the compensation matrix was then applied to the sample raw image files to create compensated image files (.cif) which were then analyzed.

### Statistical Analyses

Descriptive statistical analyses were conducted using GraphPad Prism. A two-way ANOVA was used whenever an experimental plan included a comparison between more than two groups followed by Bonferroni post-tests between groups of interest. Student t-tests were utilized wherever comparisons between only two groups were made. Details regarding sample size, the specific statistical test employed, and corresponding p-values for each figure, are provided in the figure legends, the results section, and/or the Methodology section. All bar graphs show mean +/− standard deviation.

### Single Cell Analyses

Epithelial cell data from the Kumar et al. Human Breast Cell Atlas dataset was acquired from cellxgene (‘scRNA-seq data - epithelial cells’ https://cellxgene.cziscience.com/collections/4195ab4c-20bd-4cd3-8b3d-65601277e731). Basal cells were subsetted based on ‘broad_cell_type’ metadata annotations. Progesterone receptor (PGR) status was assigned based on cells expressing greater than mean PGR expression, denoted as PGR negative (PGR-) or PGR positive (PGR+). The proportion of PGR+ cells per sample was calculated using the propeller function within the Speckle R package (v1.2.0, Phipson et al. 2022 https://doi.org/10.1093/bioinformatics/btac582). Single Cell Pathway Analysis (SCPA v1.5.0, Bibby et al. 2022 https://doi.org/10.1016/j.celrep.2022.111697) was used for pathway analysis using Reactome pathways from msigdbr (7.5.1). Predicted pathway alterations between PGR− and PGR+ basal cells with adjusted p-value <0.01 and q-value >1.5 were visualized.

### Proteomic sample preparation

DROPPS methods was used for proteomic analysis.^28^ Cells were deposited into wells of Teflon-coated slides (Tekdon, template available upon request) by pipette or directly by FACS. The solution accompanying the cells was allowed to evaporate at RT in a biosafety cabinet before storing slides at −80°C until further processing. For lysis and protein reduction, 2.5 μL of buffer consisting of 30% (v/v) Invitrosol (ThermoFisher, MS10007), 15% (v/v) acetonitrile, 0.06% (w/v) n-dodecyl-β-D-maltoside (Sigma, 850520P), 5 mM Tris(2-carboxyethyl)phosphine, and 100 mM ammonium bicarbonate in HPLC grade water. Slides were placed in a 37°C humidity chamber and allowed to incubate for 30 min. For alkylation and protein digestion, 1 μL of buffer containing 35 mM iodoacetamide and Trypsin/Lys-C (Promega, V5072) was added to each sample. Samples were allowed to digest for 2 h in a 37°C humidity chamber. Digestion was quenched by bringing samples to 0.1% (v/v) formic acid with 8 μL of 0.14% (v/v) formic acid in 37°C HPLC grade water and then samples were transferred to 96-well plate. The wells were washed with an additional 8 μL of 0.1% (v/v) formic acid and then the plate was transferred to EASY-nLC™ 1000 System chilled to 7°C.

### Mass spectrometry data acquisition

LC-MS/MS analysis was performed on an Orbitrap Fusion MS (ThermoFisher) coupled to EASY-nLC™ 1000 System (ThermoFisher). Peptides were washed on pre-column (Acclaim™ PepMap™ 100 C18, ThermoFisher) with 60 μL of mobile phase A (0.1% FA in HPLC grade water) at 3 μL/min separated using a 50 cm EASY-Spray column (ES803, ThermoFisher) ramping mobile phase B (0.1% FA in HPLC grade acetonitrile) from 0% to 5% in 2 min, 5% to 27% in 160 min, 27% to 60% in 40 min interfaced online using an EASY-Spray™ source (ThermoFisher). The Orbitrap Fusion MS was operated in data dependent acquisition mode using a 2.5 s cycle at a full MS resolution of 240,000 with a full scan range of 350-1550 *m/z* with RF Lens at 60%, full MS AGC at 200%, and maximum inject time at 40 ms. MS/MS scans were recorded in the ion trap with 1.2 Th isolation window, 100 ms maximum injection time, with a scan range of 200-1400 *m/z* using Normal scan rate. Ions for MS/MS were selected using monoisotopic peak detection, intensity threshold of 1,000, positive charge states of 2-5, 40 s dynamic exclusion, and then fragmented using HCD with 31% NCE.

### Mass spectrometry raw data analysis

Raw files were analyzed using FragPipe (v.20.0) using MSFragger^55,56^ (v.3.8) to search against a mouse (Uniprot, 25,474 sequences, accessed 2021-06-03) proteomes – canonical plus isoforms. Default settings for LFQ workflow^57,58^ were used using IonQuant^59^ v.1.9.8 and Philosopher^60^ v.5.0.0 with the following modifications: Precursor and fragment mass tolerance were specified at −50 to 50 ppm and 0.15 Da, respectively; parameter optimization was disabled; Pyro-Glu or loss of ammonia at peptide N-terminal and alkylation of N were included as variable modifications; MaxLFQ min ions was set to 1; MBR RT tolerance was set to 1 min, and MBR top runs was set to 20.

### Mass spectrometry statistical analysis

All analysis was performed using R programming language (v.4.2.2) with Tidyverse pacakge (tidyverse_1.3.2) unless otherwise specified. The “MaxLFQ Intensity” columns were extracted from the “combined_protein.tsv” output file from FragPipe. Proteins were filtered out if the Uniprot Accession ID matched one from the MaxQuant contaminants list unless they belonged to a set of IDs being used for comparison. Proteins were filtered for presence in > 50% of samples and subsequently imputed with random forest algorithm using the MissForest package (missForest_1.5). Canonical compartment marker^30^ heatmap generation was clustered using Pretty Heatmap package (pheatmap_1.0.12) using Euclidean distance for samples, and Ward D linkage. Differential expression was calculated using the Estimated Marginal Means package (emmeans_1.8.4-1) to calculate group means from a linear mixed model fit with compartment and PR as fixed effects with interaction and allowing for different intercept per mouse. GSEA was applied using a preranked list of Gene Symbols sorted based on estimated fold changes against the Mouse Reactome, Cancer Hallmarks, and Gene Ontology Biological Processes gene sets with minimum and maximum sizes of 25 and 200, respectively. Outputs from GSEA were used as inputs to Cytoscape (v.3.9.1). GSEA gene sets were visualized in EnrichmentMap (v.3.3.4) and AutoAnnotate (v.1.3.5).

### Data visualization

Unless otherwise specified, plots were generated using R programming language (v.4.2.2) with Tidyverse pacakge (tidyverse_1.3.2) with ggthemes_4.2.4, ggpubr_0.5.0, ggplot2_3.4.0, and ggbeeswarm_0.6.0 packages.

## AUTHOR CONTRIBUTIONS

Conceptualization: PT, PW, RK

Methodology: PT, PW, MW, CWM, KA, BS, RK

Formal analysis: PR, MW, CWM, KA, HB, VD

Resources: BS, SH, RC, HB, RK

Writing: PT, MW, CWM, SH, VD, PW, RK.

Visualization: PT, MW, CWM, KA.

Funding acquisition: TK, SH, RB, RK.

Supervision: RK.

## REFERENCES

1. Lydon, J. P. et al. Mice lacking progesterone receptor exhibit pleiotropic reproductive abnormalities. Genes Dev 9, 2266–78 (1995).

2. Mulac-Jericevic, B., Lydon, J. P., DeMayo, F. J. & Conneely, O. M. Defective mammary gland morphogenesis in mice lacking the progesterone receptor B isoform. Proc Natl Acad Sci U S A 100, 9744–9 (2003).

3. Beleut, M. et al. Two distinct mechanisms underlie progesterone-induced proliferation in the mammary gland. Proc Natl Acad Sci U S A 107, 2989–94 (2010).

4. Ismail, P. M., Li, J., DeMayo, F. J., O’Malley, B. W. & Lydon, J. P. A novel LacZ reporter mouse reveals complex regulation of the progesterone receptor promoter during mammary gland development. Mol Endocrinol 16, 2475–89 (2002).

5. Aupperlee, M. D. & Haslam, S. Z. Differential hormonal regulation and function of progesterone receptor isoforms in normal adult mouse mammary gland. Endocrinology 148, 2290–300 (2007).

6. Joshi, P. A. et al. Progesterone induces adult mammary stem cell expansion. Nature 465, 803–807 (2010).

7. Asselin-Labat, M.-L. et al. Control of mammary stem cell function by steroid hormone signalling. Nature 465, 798–802 (2010).

8. Graham, J. D. et al. DNA replication licensing and progenitor numbers are increased by progesterone in normal human breast. Endocrinology 150, 3318–26 (2009).

9. Hilton, H. N. et al. Progesterone stimulates progenitor cells in normal human breast and breast cancer cells. Breast Cancer Res Treat 143, 423–433 (2014).

10. Fata, J. E., Chaudhary, V. & Khokha, R. Cellular Turnover in the Mammary Gland Is Correlated with Systemic Levels of Progesterone and Not 17β-Estradiol During the Estrous Cycle1. Biol Reprod 65, 680–688 (2001).

11. Brisken, C. et al. A paracrine role for the epithelial progesterone receptor in mammary gland development. Proc Natl Acad Sci U S A 95, 5076–81 (1998).

12. Stingl, J. et al. Purification and unique properties of mammary epithelial stem cells. Nature 439, 993–7 (2006).

13. Asselin-Labat, M.-L. et al. Steroid hormone receptor status of mouse mammary stem cells. J Natl Cancer Inst 98, 1011–4 (2006).

14. Shackleton, M. et al. Generation of a functional mammary gland from a single stem cell. Nature 439, 84–8 (2006).

15. Brisken, C. Progesterone signalling in breast cancer: a neglected hormone coming into the limelight. Nat Rev Cancer 13, 385–96 (2013).

16. Lydon, J. P., Ge, G., Kittrell, F. S., Medina, D. & O’Malley, B. W. Murine mammary gland carcinogenesis is critically dependent on progesterone receptor function. Cancer Res 59, 4276–84 (1999).

17. Lee, O., Choi, M.-R., Christov, K., Ivancic, D. & Khan, S. A. Progesterone receptor antagonism inhibits progestogen-related carcinogenesis and suppresses tumor cell proliferation. Cancer Lett 376, 310–317 (2016).

18. Mørch, L. S. et al. Contemporary Hormonal Contraception and the Risk of Breast Cancer. N Engl J Med 377, 2228–2239 (2017).

19. Collaborative Group on Hormonal Factors in Breast Cancer. Type and timing of menopausal hormone therapy and breast cancer risk: individual participant meta-analysis of the worldwide epidemiological evidence. Lancet 394, 1159–1168 (2019).

20. Chlebowski, R. T. et al. Randomized trials of estrogen-alone and breast cancer incidence: a meta-analysis. Breast Cancer Res Treat 206, 177–184 (2024).

21. Sigl, V. et al. RANKL/RANK control Brca1 mutation-driven mammary tumors. Cell Res 26, 761–74 (2016).

22. Schramek, D. et al. Osteoclast differentiation factor RANKL controls development of progestin-driven mammary cancer. Nature 468, 98–102 (2010).

23. Poole, A. J. et al. Prevention of Brca1-mediated mammary tumorigenesis in mice by a progesterone antagonist. Science 314, 1467–70 (2006).

24. Gonzalez-Suarez, E. et al. RANK ligand mediates progestin-induced mammary epithelial proliferation and carcinogenesis. Nature 468, 103–7 (2010).

25. Ranjan, M. et al. Progesterone receptor antagonists reverse stem cell expansion and the paracrine effectors of progesterone action in the mouse mammary gland. Breast Cancer Research 23, 78 (2021).

26. Van Keymeulen, A. et al. Lineage-Restricted Mammary Stem Cells Sustain the Development, Homeostasis, and Regeneration of the Estrogen Receptor Positive Lineage. Cell Rep 20, 1525–1532 (2017).

27. Shehata, M. et al. Phenotypic and functional characterization of the luminal cell hierarchy of the mammary gland. Breast Cancer Res 14, R134 (2012).

28. Waas, M. et al. Droplet-based proteomics reveals CD36 as a marker for progenitors in mammary basal epithelium. Cell reports methods 4, 100741 (2024).

29. Mahendralingam, M. J. et al. Mammary epithelial cells have lineage-rooted metabolic identities. Nat Metab 3, 665–681 (2021).

30. Casey, A. E. et al. Mammary molecular portraits reveal lineage-specific features and progenitor cell vulnerabilities. J Cell Biol 217, 2951–2974 (2018).

31. Pal, B. et al. Construction of developmental lineage relationships in the mouse mammary gland by single-cell RNA profiling. Nat Commun 8, 1–13 (2017).

32. Bach, K. et al. Differentiation dynamics of mammary epithelial cells revealed by single-cell RNA sequencing. Nat Commun 8, (2017).

33. Gray, G. K. et al. A human breast atlas integrating single-cell proteomics and transcriptomics. Dev Cell 57, 1400–1420.e7 (2022).

34. Kumar, T. et al. A spatially resolved single-cell genomic atlas of the adult human breast. Nature 620, 181–191 (2023).

35. Deng, G., Lu, Y., Zlotnikov, G., Thor, A. D. & Smith, H. S. Loss of heterozygosity in normal tissue adjacent to breast carcinomas. Science 274, 2057–9 (1996).

36. Nguyen, Q. H. et al. Profiling human breast epithelial cells using single cell RNA sequencing identifies cell diversity. Nat Commun 9, 2028 (2018).

37. Rios, A. C., Fu, N. Y., Lindeman, G. J. & Visvader, J. E. In situ identification of bipotent stem cells in the mammary gland. Nature 506, 322–7 (2014).

38. Wang, D. et al. Identification of multipotent mammary stem cells by protein C receptor expression. Nature 517, 81–4 (2015).

39. Van Keymeulen, A. et al. Distinct stem cells contribute to mammary gland development and maintenance. Nature 479, 189–193 (2011).

40. Wuidart, A. et al. Quantitative lineage tracing strategies to resolve multipotency in tissue-specific stem cells. Genes Dev 30, 1261–77 (2016).

41. Wang, C., Christin, J. R., Oktay, M. H. & Guo, W. Lineage-Biased Stem Cells Maintain Estrogen-Receptor-Positive and -Negative Mouse Mammary Luminal Lineages. Cell Rep 18, 2825–2835 (2017).

42. Hilton, H. N. et al. Progesterone and estrogen receptors segregate into different cell subpopulations in the normal human breast. Mol Cell Endocrinol 361, 191–201 (2012).

43. Raouf, A. et al. Transcriptome analysis of the normal human mammary cell commitment and differentiation process. Cell Stem Cell 3, 109–18 (2008).

44. Smith, G. H. Label-retaining epithelial cells in mouse mammary gland divide asymmetrically and retain their template DNA strands. Development 132, 681–7 (2005).

45. Clarke, R. B. et al. A putative human breast stem cell population is enriched for steroid receptor-positive cells. Dev Biol 277, 443–456 (2005).

46. Booth, B. W. & Smith, G. H. Estrogen receptor-α and progesterone receptor are expressed in label-retaining mammary epithelial cells that divide asymmetrically and retain their template DNA strands. Breast Cancer Research 8, R49 (2006).

47. Masciari, S. et al. Breast cancer phenotype in women with TP53 germline mutations: a Li-Fraumeni syndrome consortium effort. Breast Cancer Res Treat 133, 1125–1130 (2012).

48. Garbe, J. C. et al. Accumulation of multipotent progenitors with a basal differentiation bias during aging of human mammary epithelia. Cancer Res 72, 3687–701 (2012).

49. Keller, P. J. et al. Defining the cellular precursors to human breast cancer. Proc Natl Acad Sci U S A 109, 2772–2777 (2012).

50. Fata, J. E. et al. The Osteoclast Differentiation Factor Osteoprotegerin-Ligand Is Essential for Mammary Gland Development. Cell 103, 41–50 (2000).

51. Sleeman, K. E. et al. Dissociation of estrogen receptor expression and in vivo stem cell activity in the mammary gland. J Cell Biol 176, 19–26 (2007).

52. Chang, T. H.-T. et al. New insights into lineage restriction of mammary gland epithelium using parity-identified mammary epithelial cells. Breast Cancer Res 16, R1 (2014).

53. Rodilla, V. et al. Luminal progenitors restrict their lineage potential during mammary gland development. PLoS Biol 13, e1002069 (2015).

54. Bartlett, T. E. et al. Antiprogestins reduce epigenetic field cancerization in breast tissue of young healthy women. Genome Med 14, 64 (2022).

55. Kong, A. T., Leprevost, F. V, Avtonomov, D. M., Mellacheruvu, D. & Nesvizhskii, A. I. MSFragger: ultrafast and comprehensive peptide identification in mass spectrometry– based proteomics. Nat Methods 14, 513–520 (2017).

56. Teo, G. C., Polasky, D. A., Yu, F. & Nesvizhskii, A. I. Fast Deisotoping Algorithm and Its Implementation in the MSFragger Search Engine. J Proteome Res 20, 498–505 (2021).

57. Nesvizhskii, A. I., Keller, A., Kolker, E. & Aebersold, R. A Statistical Model for Identifying Proteins by Tandem Mass Spectrometry. Anal Chem 75, 4646–4658 (2003).

58. Käll, L., Canterbury, J. D., Weston, J., Noble, W. S. & MacCoss, M. J. Semi-supervised learning for peptide identification from shotgun proteomics datasets. Nat Methods 4, 923–925 (2007).

59. Yu, F., Haynes, S. E. & Nesvizhskii, A. I. IonQuant Enables Accurate and Sensitive Label-Free Quantification With FDR-Controlled Match-Between-Runs. Molecular & Cellular Proteomics 20, 100077 (2021).

60. da Veiga Leprevost, F., et al. Philosopher: a versatile toolkit for shotgun proteomics data analysis. Nat Methods 17, 869–870 (2020).

