## Supplemental Figures for "PR+ progenitors contribute to all mammary epithelial lineages"

#### SUPPLEMENTALS & SUPPLEMENTAL FIGURE LEGENDS

**Figure S1 Flow gating strategy**

(A) Basic gating strategy used for all flow cytometric analyses of mammary cells.

(B) Gating strategy used for flow cytometric analyses of cells from the $PR^{LacZ}$ mouse model with the FDG/LacZ flow cytometry kit, where LacZ signal serves as a reporter for both PRA and PRB.

#### Figure S1

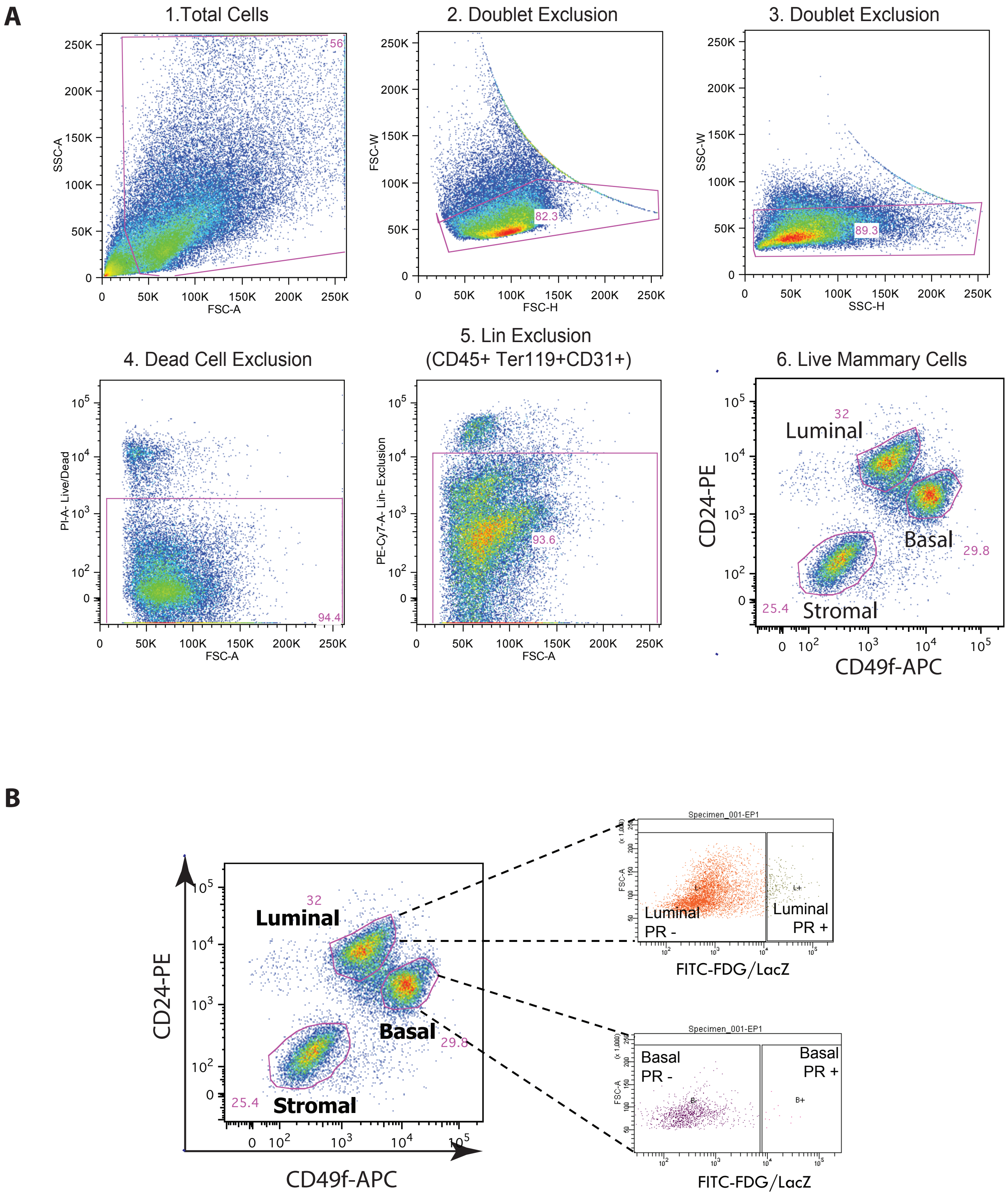
A
1.Total Cells
2. Doublet Exclusion
3. Doublet Exclusion
4. Dead Cell Exclusion
5. Lin Exclusion
(CD45+ Ter119+CD31+)
6. Live Mammary Cells
SSC-A
FSC-A
FSC-W
FSC-H
SSC-W
SSC-H
PI-A- Live/Dead
PE-Cy7-A- Lin- Exclusion
CD24-PE
CD49f-APC
Luminal
Basal
Stromal
56
82.3
89.3
94.4
93.6
32
29.8
25.4
B
Specimen_001-EP1
FSC-A
Luminal
PR -
Luminal
PR +
FITC-FDG/LacZ
Basal
PR -
Basal
PR +

**Figure S2 Basal PR+ cells**

(A) Dual immunofluorescence of tissue sections from WT mice showing localization of the basal marker Keratin 14 with total PR, PRA and PRB isoform-specific antibodies (top; scale bar = 20 $\mu$m). Dual immunofluorescence of tissue sections from PR LacZ mice shows cellular co-localization of PR and β-galactosidase (bottom; scale bar = 50 $\mu$m).

(B) Immunofluorescent staining of PR and lineage markers in luminal and basal cells that were FACS-sorted based on the FDG/LacZ-signal. (Scale bar = 20 $\mu$m).

(C) Colony forming capacity relative to the number of cells plated of FACS-sorted PR LacZ+ and PR LacZ- luminal and basal cells from E and EP treated PR LacZ littermate wildtype control mice and PRKO LacZ mice.

(D) Flow plots showing the enrichment of PR LacZ+ cells within the MRU basal fraction in EP treated PR LacZ mice.

(E) qRT-PCR results from RNA extracted from FACS-sorted cells from the basal (B), MRU, and myoepithelial (MYO) fractions from wild type E and EP treated mice. Expression levels were normalized to β-actin and the fold-change calculated relative to E treated basal cells.

### Figure S2

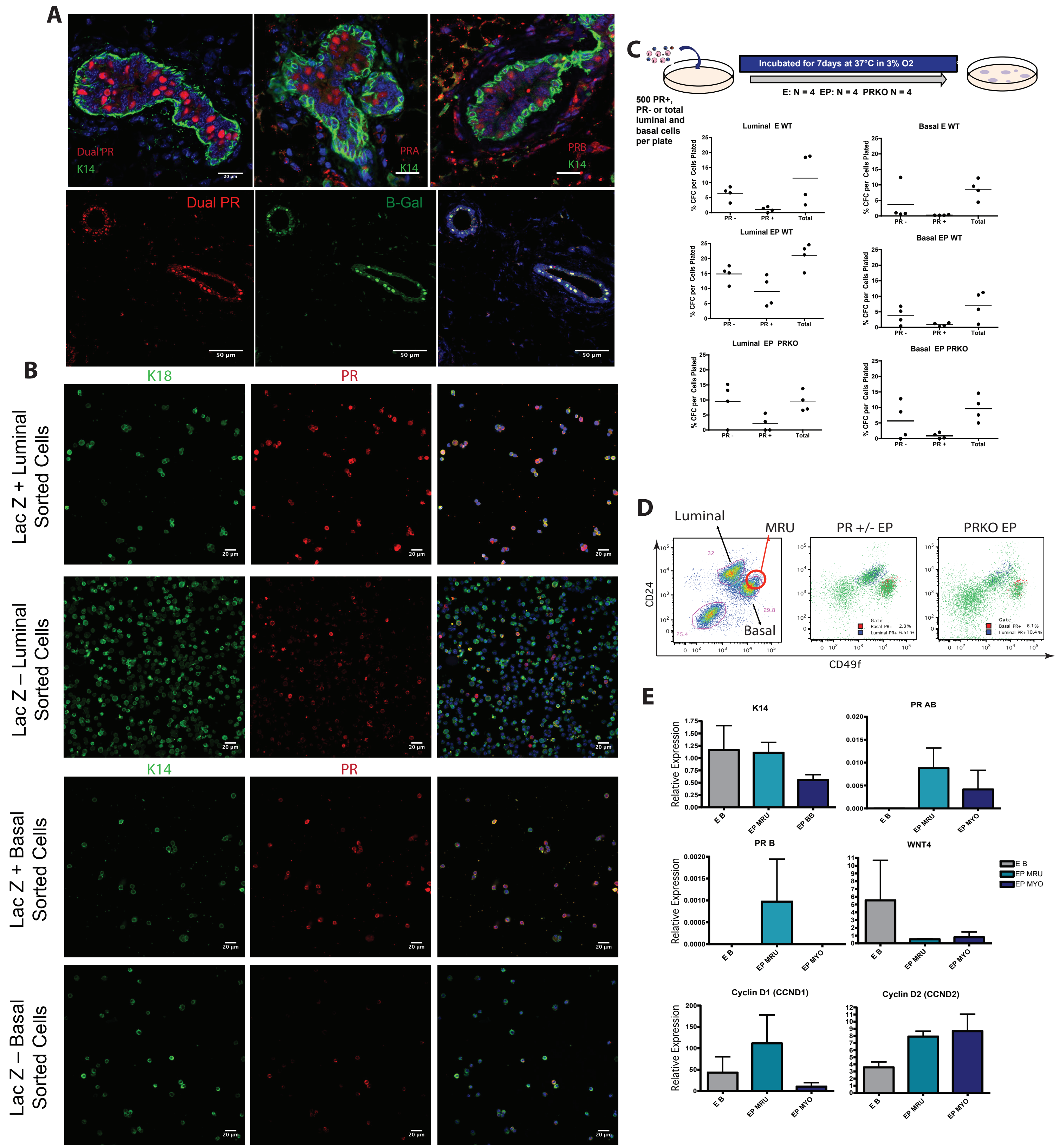
A
Dual PR
K14
PRA
K14
PRB
K14
20 µm
Dual PR
B-Gal
50 µm
B
K18
PR
Lac Z + Luminal Sorted Cells
Lac Z – Luminal Sorted Cells
K14
PR
Lac Z + Basal Sorted Cells
Lac Z – Basal Sorted Cells
20 µm
C
500 PR+, PR- or total luminal and basal cells per plate
Incubated for 7days at 37°C in 3% O2
E: N = 4 EP: N = 4 PRKO N = 4
Luminal E WT
Basal E WT
Luminal EP WT
Basal EP WT
Luminal EP PRKO
Basal EP PRKO
% CFC per Cells Plated
PR -
PR +
Total
D
Luminal
MRU
Basal
PR +/- EP
PRKO EP
CD24
CD49f
E
K14
PR AB
PR B
WNT4
Cyclin D1 (CCND1)
Cyclin D2 (CCND2)
Relative Expression
E B
EP MRU
EP MYO
EP BB

**Figure S3. Low input proteomic profiling of PR+ basal cells**

(A) PCA component analysis for PR status and cell lineage variables.

(B) The number of proteins with significant differences between the four different cell types.

(C,D) The Spearman's correlations between (C) the estimated mean log intensities of luminal and basal cells with or without PR and (D) the estimated fold changes between the four cell types.

(E) A dot plot depicting the most significant pathways between basal and luminal lineages within PR+ and PR- subsets.

(F) The GSEA NSE values for significant pathways for basal-luminal comparisons are plotted for the PR- and PR+ subsets. The Spearman's correlation for the NES values is shown.

### Figure S3

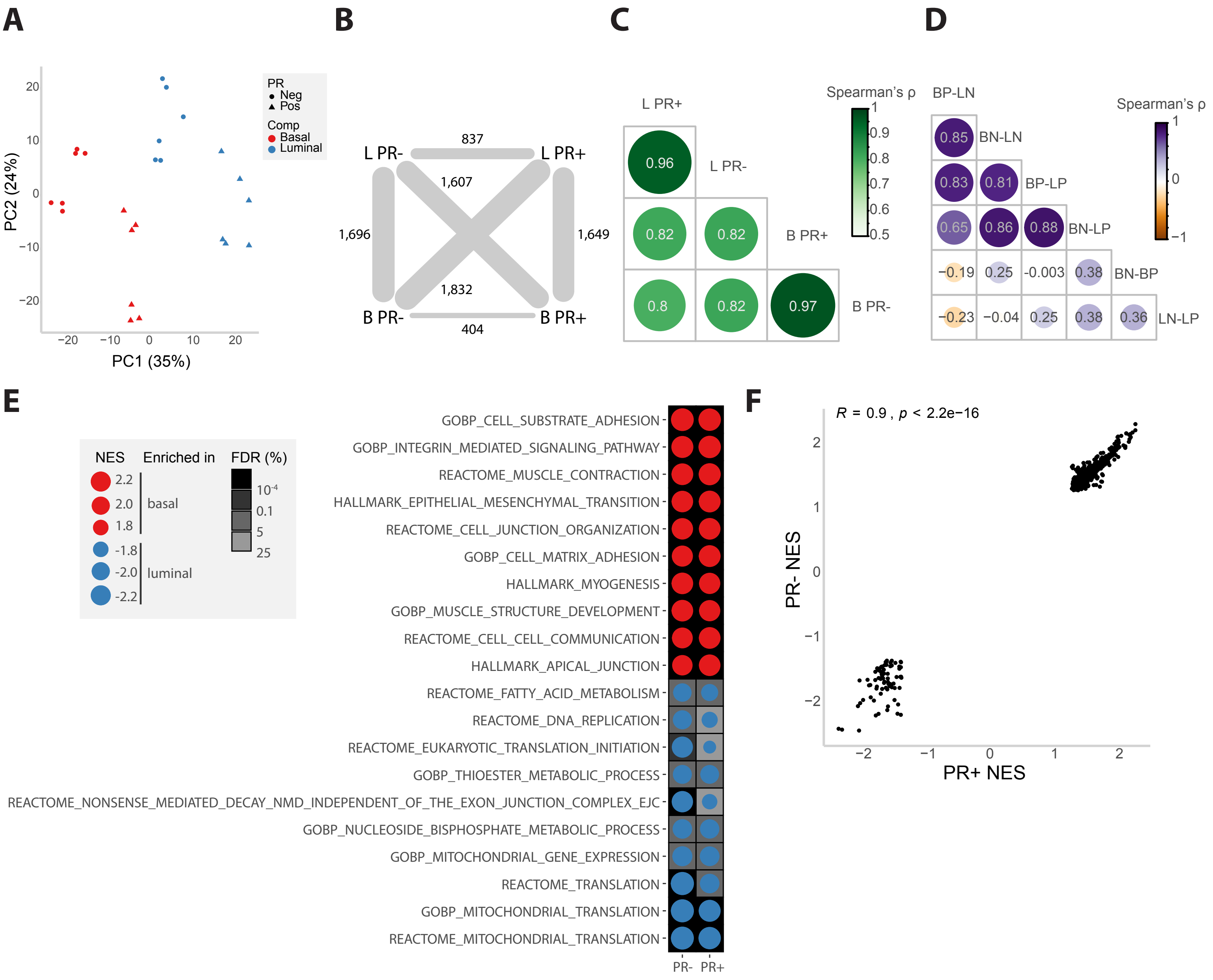
A
PC2 (24%)
PC1 (35%)
PR
Neg
Pos
Comp
Basal
Luminal
B
L PR-
L PR+
B PR-
B PR+
837
1,607
1,696
1,649
1,832
404
C
Spearman's ρ
L PR+
L PR-
B PR+
B PR-
0.96
0.82
0.82
0.8
0.82
0.97
D
Spearman's ρ
BP-LN
BN-LN
BP-LP
BN-LP
BN-BP
LN-LP
0.85
0.83
0.81
0.65
0.86
0.88
−0.19
0.25
-0.003
0.38
−0.23
−0.04
0.25
0.38
0.36
E
NES
Enriched in
FDR (%)
basal
luminal
GOBP_CELL_SUBSTRATE_ADHESION
GOBP_INTEGRIN_MEDIATED_SIGNALING_PATHWAY
REACTOME_MUSCLE_CONTRACTION
HALLMARK_EPITHELIAL_MESENCHYMAL_TRANSITION
REACTOME_CELL_JUNCTION_ORGANIZATION
GOBP_CELL_MATRIX_ADHESION
HALLMARK_MYOGENESIS
GOBP_MUSCLE_STRUCTURE_DEVELOPMENT
REACTOME_CELL_CELL_COMMUNICATION
HALLMARK_APICAL_JUNCTION
REACTOME_FATTY_ACID_METABOLISM
REACTOME_DNA_REPLICATION
REACTOME_EUKARYOTIC_TRANSLATION_INITIATION
GOBP_THIOESTER_METABOLIC_PROCESS
REACTOME_NONSENSE_MEDIATED_DECAY_NMD_INDEPENDENT_OF_THE_EXON_JUNCTION_COMPLEX_EJC
GOBP_NUCLEOSIDE_BISPHOSPHATE_METABOLIC_PROCESS
GOBP_MITOCHONDRIAL_GENE_EXPRESSION
REACTOME_TRANSLATION
GOBP_MITOCHONDRIAL_TRANSLATION
REACTOME_MITOCHONDRIAL_TRANSLATION
PR-
PR+
F
R = 0.9 , p < 2.2e−16
PR- NES
PR+ NES

**Figure S4. PGR interrogation in public human breast single-cell RNAseq data.**

(A, B) Proportion of PGR+ basal cells (A) and PGR+ luminal hormone sensing cells (B) per pre-menopausal sample in the Kumar et al. 2023 Human Breast Cell Atlas single-cell RNAseq dataset.

(C) Estimation of gene expression density for select mammary lineage marker genes in the Kumar et al. 2023 Human Breast Cell Atlas single-cell RNAseq dataset.

(D) Reactome pathway analysis of PGR+ versus PGR- luminal hormone sensing (top), PR+ luminal hormone sensing versus PGR+ basal (middle) and PGR- luminal hormone sensing vs. PGR- basal (bottom) in the Kumar et al. 2023 Human Breast Cell Atlas single-cell RNAseq dataset.

**Table 1-1 qRT-PCR Primer List**

| Applied Biosystems Gene Expression Assay Catalogue Numbers | |
|---|---|
| Gene Name | Catalogue number |
| *Keratin 18* | Mm01601706_g1 |
| *Keratin 14* | Mm00516876_m1 |
| *Rankl* | Mm01313944_g1 |
| *Rank* | Mm01286484_m1 |
| *Wnt 4* | Mm01194003_m1 |
| *Cyclin D1* | Rn00596851_g1 |
| *Cyclin D2* | Mm03053712_s1 |
| *$\beta$-actin (ACTB)* | Mm01205647_g1 |
| Custom Primers /Probes | Sequence |
| *PR-B* | |
| Probe | 5’ - CTTCCAAGGGAGCCAGCACTCGG - 3' |
| Forward | 5’ - CGCAGGTTCTCCACACGTC - 3’ |
| Reverse | 5’ - GATCGGTATAGGCGAGACTACAGAC - 3’ |
| *PR-AB* | |
| Probe | 5’ - CACGCCATAGTGACAGCCAGATGCTT - 3’ |
| Forward | 5’ - CACAGTATGGCTTTGATTCCTTACCTC - 3’ |
| Reverse | 5’ - TGCCCTCTTAAAGAAGACCTTGC -3’ |

### Figure S4

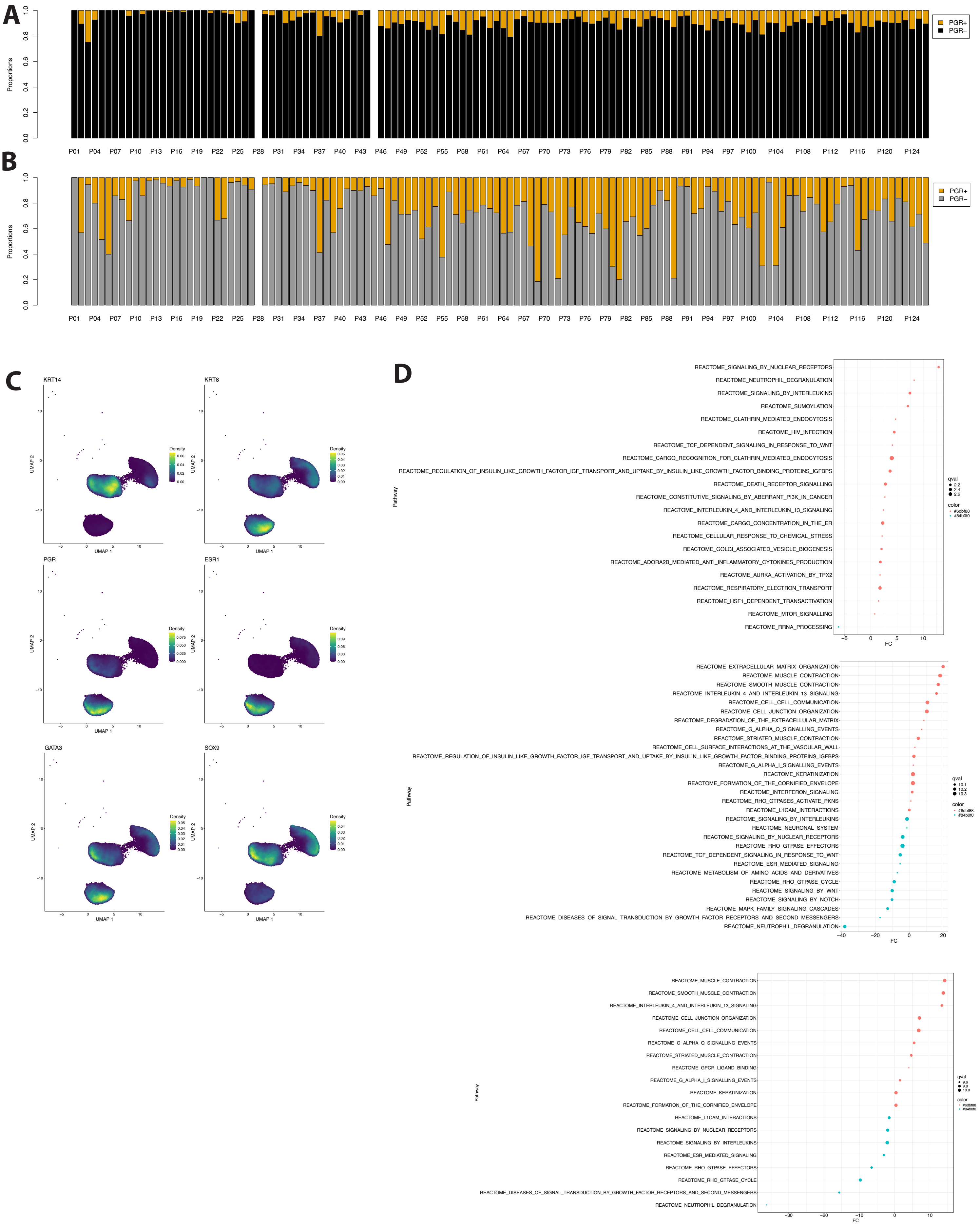
A
B
C
D
Proportions
PGR+
PGR−
P01 P04 P07 P10 P13 P16 P19 P22 P25 P28 P31 P34 P37 P40 P43 P46 P49 P52 P55 P58 P61 P64 P67 P70 P73 P76 P79 P82 P85 P88 P91 P94 P97 P100 P104 P108 P112 P116 P120 P124
KRT14
KRT8
PGR
ESR1
GATA3
SOX9
Density
UMAP 1
UMAP 2
Pathway
FC
qval
color
#6dbf88
#84b0f0
REACTOME_SIGNALING_BY_NUCLEAR_RECEPTORS
REACTOME_NEUTROPHIL_DEGRANULATION
REACTOME_SIGNALING_BY_INTERLEUKINS
REACTOME_SUMOYLATION
REACTOME_CLATHRIN_MEDIATED_ENDOCYTOSIS
REACTOME_HIV_INFECTION
REACTOME_TCF_DEPENDENT_SIGNALING_IN_RESPONSE_TO_WNT
REACTOME_CARGO_RECOGNITION_FOR_CLATHRIN_MEDIATED_ENDOCYTOSIS
REACTOME_REGULATION_OF_INSULIN_LIKE_GROWTH_FACTOR_IGF_TRANSPORT_AND_UPTAKE_BY_INSULIN_LIKE_GROWTH_FACTOR_BINDING_PROTEINS_IGFBPS
REACTOME_DEATH_RECEPTOR_SIGNALLING
REACTOME_CONSTITUTIVE_SIGNALING_BY_ABERRANT_PI3K_IN_CANCER
REACTOME_INTERLEUKIN_4_AND_INTERLEUKIN_13_SIGNALING
REACTOME_CARGO_CONCENTRATION_IN_THE_ER
REACTOME_CELLULAR_RESPONSE_TO_CHEMICAL_STRESS
REACTOME_GOLGI_ASSOCIATED_VESICLE_BIOGENESIS
REACTOME_ADORA2B_MEDIATED_ANTI_INFLAMMATORY_CYTOKINES_PRODUCTION
REACTOME_AURKA_ACTIVATION_BY_TPX2
REACTOME_RESPIRATORY_ELECTRON_TRANSPORT
REACTOME_HSF1_DEPENDENT_TRANSACTIVATION
REACTOME_MTOR_SIGNALLING
REACTOME_RRNA_PROCESSING
REACTOME_EXTRACELLULAR_MATRIX_ORGANIZATION
REACTOME_MUSCLE_CONTRACTION
REACTOME_SMOOTH_MUSCLE_CONTRACTION
REACTOME_INTERLEUKIN_4_AND_INTERLEUKIN_13_SIGNALING
REACTOME_CELL_CELL_COMMUNICATION
REACTOME_CELL_JUNCTION_ORGANIZATION
REACTOME_DEGRADATION_OF_THE_EXTRACELLULAR_MATRIX
REACTOME_G_ALPHA_Q_SIGNALLING_EVENTS
REACTOME_STRIATED_MUSCLE_CONTRACTION
REACTOME_CELL_SURFACE_INTERACTIONS_AT_THE_VASCULAR_WALL
REACTOME_REGULATION_OF_INSULIN_LIKE_GROWTH_FACTOR_IGF_TRANSPORT_AND_UPTAKE_BY_INSULIN_LIKE_GROWTH_FACTOR_BINDING_PROTEINS_IGFBPS
REACTOME_G_ALPHA_I_SIGNALLING_EVENTS
REACTOME_KERATINIZATION
REACTOME_FORMATION_OF_THE_CORNIFIED_ENVELOPE
REACTOME_INTERFERON_SIGNALING
REACTOME_RHO_GTPASES_ACTIVATE_PKNS
REACTOME_L1CAM_INTERACTIONS
REACTOME_SIGNALING_BY_INTERLEUKINS
REACTOME_NEURONAL_SYSTEM
REACTOME_SIGNALING_BY_NUCLEAR_RECEPTORS
REACTOME_RHO_GTPASE_EFFECTORS
REACTOME_TCF_DEPENDENT_SIGNALING_IN_RESPONSE_TO_WNT
REACTOME_ESR_MEDIATED_SIGNALING
REACTOME_METABOLISM_OF_AMINO_ACIDS_AND_DERIVATIVES
REACTOME_RHO_GTPASE_CYCLE
REACTOME_SIGNALING_BY_WNT
REACTOME_SIGNALING_BY_NOTCH
REACTOME_MAPK_FAMILY_SIGNALING_CASCADES
REACTOME_DISEASES_OF_SIGNAL_TRANSDUCTION_BY_GROWTH_FACTOR_RECEPTORS_AND_SECOND_MESSENGERS
REACTOME_NEUTROPHIL_DEGRANULATION
REACTOME_MUSCLE_CONTRACTION
REACTOME_SMOOTH_MUSCLE_CONTRACTION
REACTOME_INTERLEUKIN_4_AND_INTERLEUKIN_13_SIGNALING
REACTOME_CELL_JUNCTION_ORGANIZATION
REACTOME_CELL_CELL_COMMUNICATION
REACTOME_G_ALPHA_Q_SIGNALLING_EVENTS
REACTOME_STRIATED_MUSCLE_CONTRACTION
REACTOME_GPCR_LIGAND_BINDING
REACTOME_G_ALPHA_I_SIGNALLING_EVENTS
REACTOME_KERATINIZATION
REACTOME_FORMATION_OF_THE_CORNIFIED_ENVELOPE
REACTOME_L1CAM_INTERACTIONS
REACTOME_SIGNALING_BY_NUCLEAR_RECEPTORS
REACTOME_SIGNALING_BY_INTERLEUKINS
REACTOME_ESR_MEDIATED_SIGNALING
REACTOME_RHO_GTPASE_EFFECTORS
REACTOME_RHO_GTPASE_CYCLE
REACTOME_DISEASES_OF_SIGNAL_TRANSDUCTION_BY_GROWTH_FACTOR_RECEPTORS_AND_SECOND_MESSENGERS
REACTOME_NEUTROPHIL_DEGRANULATION
